# Molecular simulations reveal a mechanism for enhanced allosteric coupling between voltage-sensor and pore domains in KCNQ1 explaining its activation by ML277

**DOI:** 10.1101/2022.05.31.494217

**Authors:** Georg Kuenze, Carlos G. Vanoye, Mason C. Wilkinson, Reshma R. Desai, Sneha Adusumilli, Charles R. Sanders, Alfred L. George, Jens Meiler

## Abstract

The voltage-gated potassium channel KCNQ1 (K_V_7.1) is important for the repolarizing phase of the cardiac action potential. Activators of KCNQ1 may provide a strategy for the pharmacological treatment of congenital long QT syndrome, a genetic disorder caused by pathogenic variants in KCNQ1 that promote arrhythmia susceptibility and elevate risk for sudden cardiac death. The small-molecule agonist ML277 recovers function of mutant KCNQ1 channels in human induced pluripotent stem cell-derived cardiomyocytes and could represent a starting point for drug development. Here we investigated ML277 mode of action by developing a molecular model of the KCNQ1-ML277 interaction corroborated by experimental and computational analyses. Ligand docking and molecular dynamics simulation demonstrated that ML277 binds to the interface between the voltage sensor and pore domains in KCNQ1. Model predicted binding energies for ML277 and 62 chemical analogs of ML277 correlated with EC_50_ data available for these compounds. We identified novel ML277-interacting residues on the S5 and S6 segments of KCNQ1 by performing MM/PBSA energy calculations and site-directed mutagenesis of KCNQ1 coupled to electrophysiological characterization of the generated channel mutants. Network analysis of the molecular dynamics simulations further showed that ML277 increases the allosteric coupling efficiency between residues in the voltage sensor domain and residues in the pore domain. Derivatives of ML277 that are not active on KCNQ1 fail to increase allosteric coupling efficiency in the computational simulations. Our results reveal atomic details of the ML277 modulation of KCNQ1 activation. These findings may be useful for the design of allosteric modulators of KCNQ1 and other KCNQ channels that bind at the membrane-accessible protein surface.

**Statement of Significance:** The potassium ion channel KCNQ1 contributes to the generation of electrical impulses in the heart. Heritable mutations in KCNQ1 can cause channel loss-of-function and predispose to a life-threatening cardiac arrhythmia. Small molecules that bind KCNQ1 and enhance channel function could establish a novel anti-arrhythmic drug paradigm. We used molecular simulations to investigate how a small agonist of KCNQ1 (ML277) binds to the KCNQ1 channel and increases its function. We identified amino acids that are responsible for ML277 binding and show how ML277 promotes signaling in KCNQ1 and channel opening. This work advances our understanding how KCNQ1 and possibly other potassium channels can be activated with small molecules. These data provide a framework for drug development studies.

## Introduction

KCNQ1 (K_V_7.1) is a voltage-gated potassium (K_V_) channel that complexes with KCNE auxiliary proteins to serve diverse physiological functions such as cardiac action potential repolarization, maintenance of ion homeostasis, and hormone secretion (1). A hallmark of KCNQ1 is its co-assembly with KCNE1 in cardiac tissue to generate a channel complex that is responsible for the slow delayed rectifier current (I_Ks_) (2, 3). I_Ks_ is an important driver of the repolarizing phase of the cardiac action potential. Heritable mutations in KCNQ1 and KCNE1 are responsible for ∼50% of cases of long QT syndrome (LQTS) (4) – a genetic disorder associated with susceptibility to potentially life-threatening ventricular arrhythmia. Pharmacological activation of I_Ks_ channels represents a novel strategy for the development of LQTS pharmacotherapy.

KCNQ1 adopts the typical structural organization of tetrameric K_V_ channels with a central pore domain (PD), formed by transmembrane (TM) segments S5 and S6 of all subunits, and four peripheral voltage-sensing domains (VSDs), each made up of four TM segments (S1-S4). A helical linker between S4 and S5 (S4-S5L) connects the VSD to the PD and is important for linking voltage-activated movements of the VSD to pore opening, a process termed VSD-PD coupling. Based on observations derived from the first crystal structure of a domain-swapped K_V_ channel (5) showing that S4-S5L sits on the C-terminal end of S6, it was initially suggested that S4-S5L may act as a ligand that holds the KCNQ1 channel in a closed state (6, 7). Movement of S4 upon membrane depolarization is thought to exert a lateral and upward pull on S4-S5L, allowing S6 to kink and making the channel permeable for ions. Work over recent years has contributed to a more complete mechanistic understanding of VSD-PD coupling in KCNQ1 (8–13). It was found that different protein interfaces are employed in the intermediate and fully activated state for communicating movements in the VSD to the pore (11, 12). Furthermore, it is now known that KCNQ1 modulators such as Phosphatidylinositol 4,5-bisphosphate (PIP2) (8, 9), KCNE1 (14), and calmodulin (CaM) (13) can influence VSD-PD coupling. The cryo-electron microscopy (EM) structure of KCNQ1 in complex with PIP2 (15) showed that PIP2 binds residues in the S2-S3 linker (S2-S3L), S4, and S4-S5L, placing it in a region where it can interact with both the VSD and PD and affect VSD-PD coupling. Depletion of PIP2 suppresses I_Ks_ current but does not prevent S4 movement (8, 9), suggesting that loss of PIP2 leads to uncoupling of the VSD from the PD. Recently, it was hypothesized that the role of PIP2 could also involve displacing CaM from interactions with the VSD so that the S6 C-terminus of KCNQ1 can change its conformation as is necessary for gate opening (13). Several lines of evidence have been collected showing that KCNE1 also modulates VSD-PD interactions (12, 14, 16– 18). Specifically, it prevents opening of KCNQ1 when the VSD is in the intermediate state and modifies the properties of the fully activated-open state, which could explain many specifics of the KCNQ1-KCNE1 channel (14).

In addition to auxiliary proteins and lipid cofactors, different small-molecule modulators of KCNQ1 have been reported in the literature. Zinc pyrithione (19) and L-364,373 (R-L3) (20) activate homomeric KCNQ1 channels, mefenamic acid (21), 4,4’-diisothiocyano-2,2’-stilbenedisulfonic acid (22), and rottlerin (23) activate the KCNQ1-KCNE1 complex, and phenylboronic acid (24), hexachlorophene (25), and 3-ethyl-2-hydroxy-2-cyclopenten-1-one (26) potentiate both KCNQ1 and KCNQ1-KCNE1. While some of these activators are able to shorten the action potential duration in isolated cardiac myocytes (20, 25), they have low potency and are not selective against other members of the KCNQ family or the hERG potassium channel. For instance, mefenamic acid (21) and phenylboronic acid (24) have reported EC_50_ values of only 60 µM and 1.6 mM, respectively. In contrast, ML277 ((R)-N-(4-(4-methoxyphenyl)thiazol-2-yl)-1-tosylpiperidine-2-carboxamide) is a potent small-molecule activator of KCNQ1 with a reported EC_50_ of 260 nM and more than 100-fold selectivity for KCNQ1 versus KCNQ2, KCNQ4, and hERG (27, 28). ML277 was identified by a library screen of >300,000 compounds and subsequent synthetic optimization of the initial hit (27). ML277 potentiates the current amplitude of KCNQ1 in both whole-cell and single-channel configurations, hyperpolarizes the voltage-dependence of activation V_1/2_ and slows channel deactivation (28–31). Similar to other KCNQ1 activators (e.g., R-L3 (20)), the activity of ML277 on KCNQ1-KCNE1 channels depends on the subunit stoichiometry (28, 31). The effect of ML277 on KCNQ1 channel current was found to be less with greater KCNE1 expression, with little (29) or no effect (28, 30) observed at saturating KCNE1 levels. Meanwhile, ML277 does shorten the action potential duration in guinea pig and canine ventricular myocytes (29) as well as in induced pluripotent stem cell (iPSC)-derived cardiomyocytes (28). This suggests that KCNQ1-KCNE1 channels in myocytes are not fully saturated with KCNE1 and that ML277 may be useful as a LQTS therapeutic.

Much could be learned about the structural basis of KCNQ1 gating and the nature of regulatory interactions of KCNQ1 with PIP2, KCNE3, and CaM by comparing cryo-EM structures of KCNQ1 in closed and open states and in *apo* and PIP2-or KCNE3-bound states (15, 32). However, no structure of KCNQ1 in complex with a small molecule ligand has yet been reported, and the location of binding sites of activators such as ML277 remain elusive. The lack of high-resolution structural data showing how small chemical ligands bind to KCNQ1 limits our understanding of drug activation mechanisms and hampers drug design efforts to find more potent KCNQ1 agonists. This kind of information has recently become available for KCNQ2 (33) and KCNQ4 (34, 35), two homologs of KCNQ1, by determining the structures of these channels in complex with small activators. These structures have revealed the location of intramembrane drug binding sites in KCNQ2 and KCNQ4, and suggested possible mechanisms for drug activity and selectivity. The activator ztz240 was found to bind at the center of the VSD in KCNQ2 and stabilize the activated VSD conformation (33). The antiepileptic drug retigabine (36, 37), and another neuronal KCNQ channel activator, ML213 (38), were observed to bind at the pore domain of KCNQ2 and KCNQ4 and appear to activate these channels by an allosteric modulation of the pore (33–35). Careful analysis of KCNQ channel-drug interactions has also highlighted the structural basis for drug selectivity and helped to explain the insensitivity of KCNQ1 to the effects of retigabine and ML213.

We investigated the structural basis for ML277 action on KCNQ1 by means of computational simulations. Using molecular docking, we identified an intramembrane binding site for ML277 at the interface between the KCNQ1 VSD and PD, which is a region that is well suited to modulate VSD-PD coupling. We validated this binding site model by comparing model predictions with structure-activity relationship data on ML277 and confirming specific ML277-KCNQ1 interactions by site-directed mutagenesis coupled to electrophysiology. These combined analyses revealed the structural determinants of selectivity of ML277 towards KCNQ1. Furthermore, using molecular dynamics (MD) simulations and network analysis we demonstrated that ML277 increases the efficiency of allosteric coupling between the VSD and PD. These findings offer an atomically detailed explanation for how ML277 augments KCNQ1 activity.

## Results

### Molecular docking identifies an intramembrane ML277 binding site on KCNQ1

To probe the binding mode of ML277 on KCNQ1 we employed molecular docking using RosettaLigand (39, 40). ML277 docking runs were started from multiple random positions within a side pocket of KCNQ1 that is formed between two neighboring VSDs (**Fig S1**). This side pocket harbors binding sites for PIP2 and KCNE3 (15), and possibly also KCNE1 (17, 41–44). Xu et al. (29) previously suggested that ML277 is also localized to this side pocket, however, the exact binding mode remained unclear. Therefore, we set the box size of the docking grid large enough (30Å x 30Å x 30Å) such that the search space enclosed the entire side pocket. The best-scoring ML277 docking poses clustered in a region bordered on the intracellular side by S2-S3L, S4-S5L, and the loop connecting S4 to S4-S5L, and on the intramembrane side by S4 from the same KCNQ1 subunit and S5 and S6 from the neighboring subunit (**Fig S2**). Docked poses were optimized by an additional high-resolution refinement step. The refined ML277 binding model with the best interface energy and P_Near_ score was selected as the representative model (**Figure 1)**. The P_Near_ metric evaluates the energy difference between the best-scoring model and alternative models with similar structure. It is a measure of the depth of the energy well occupied by the protein-ligand structure model.

**Figure 1:**
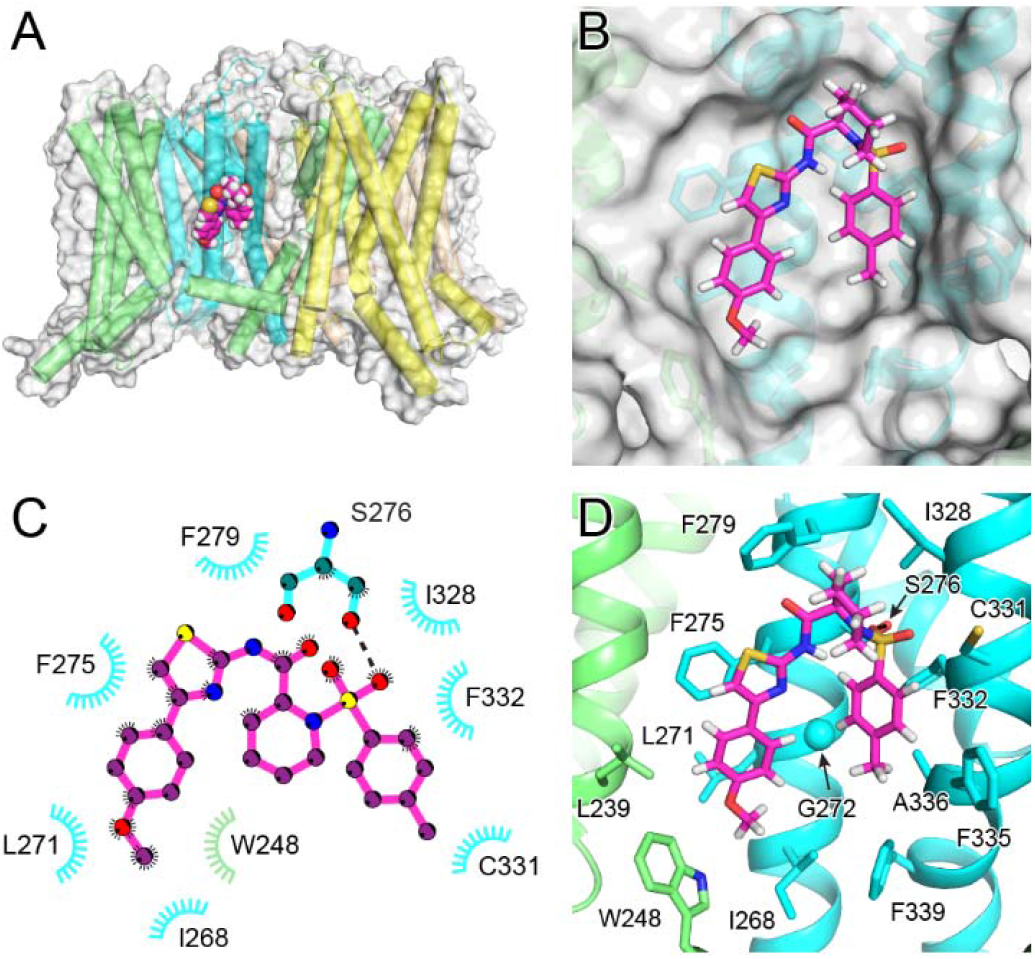
Molecular model of the KCNQ1-ML277 complex. (**A**) KCNQ1 channel is shown with cylindrical helices and in surface mode. Only the transmembrane part is shown for clarity. ML277 ligand is depicted with spheres. (**B**) Zoom-in view of ML277 binding model. ML277 fits the shape of a surface cavity in KCNQ1. (**C**) Schematic diagram of KCNQ1-ML277 interactions. (**D**) Molecular interactions between ML277 and KCNQ1 residues. KCNQ1 side chains are depicted as sticks and are labeled.

ML277 is observed to bind to a membrane-exposed surface cavity in the TM region of KCNQ1 (**Fig 1A+B**). The cavity is formed mostly by hydrophobic and aromatic residues on S5 and S6. On the cytosolic side, the cavity is bordered by S4-S5L and the C-terminal end of S4 from a neighboring KCNQ1 subunit. Control docking calculations using a different docking algorithm and scoring function as implemented in the MOE software yielded an almost identical ML277 binding pose (**Fig S3A**) that was separated from similar, alternative docking poses by a steep energy funnel (**Fig S3B**).

**Figures 1C** and **1D** depict the molecular interactions formed between KCNQ1 and ML277. ML277 contains three major functional groups: the methoxyphenyl-thiazol group, the middle piperidine-carboxamide ring, and the methylphenyl-sulfonyl group. The methoxyphenyl-thiazol group of ML277 makes hydrophobic interactions with L239 on S4, W248 on S4-S5L, and I268, L271, and F275 on S5. Among those hydrophobic residues, F275 could potentially form π-π stacking interactions with the thiazol ring of ML277. The piperidine-carboxamide moiety of ML277 interacts with F279 on S5 and I328 and C331 on S6. The 4-methylpenyl-sulfonyl group interacts mostly with residues on S6: C331, F332, F335, and A336. Furthermore, the model predicts that the position of the sulfonyl group could be stabilized by a hydrogen bond with the side chain OH of S276 on S5. Residue G272 is also notable because this small amino acid offers enough space for ML277 to fit into the surface cavity and engage in interactions with the other larger side chains. Interestingly, G272 is substituted by bulkier Cys and Val residues in KCNQ2 and KCNQ4 channels, respectively, which are insensitive to ML277. Overall, the geometric shape of ML277 and the molecular surface in KCNQ1 have high complementarity (see **Fig 1B**).

### Correlation of ML277 docking scores with ligand quantitative structure-activity data

To assess the accuracy of the KCNQ1-ML277 binding model, we compared protein-ligand interface score (I_sc) and P_Near_ metric with available SAR data for ML277. The interface score is a prediction of the protein-ligand interaction energy and the P_Near_ value estimates the probability of the docked model to represent the bound state. Ideally, these values report on changes in ligand binding affinity or protein function owing to a chemical modification of the ligand. Previously, Mattmann et al. (27) determined the activities (EC_50_) of ML277 and of 62 chemical analogs of ML277 carrying modifications in either the sulfonamide group, the internal ring system, or the terminal amide portion of the molecule. We docked each ML277 analog into the identified binding site and determined I_sc and P_Near_ values. We expected that the activities of ML277 and ML277 derivatives and their respective I_sc and P_Near_ values should be correlated if the KCNQ1-ML277 binding model was accurate. Indeed, we found a clear ranking of ligand EC_50_ values with P_Near_ (Spearman’s R = -0.286, p = 0.023) and I_sc (Spearman’s R = 0.291, p = 0.021) as shown in **Figures 2A** and **2B**, respectively.

**Figure 2:**
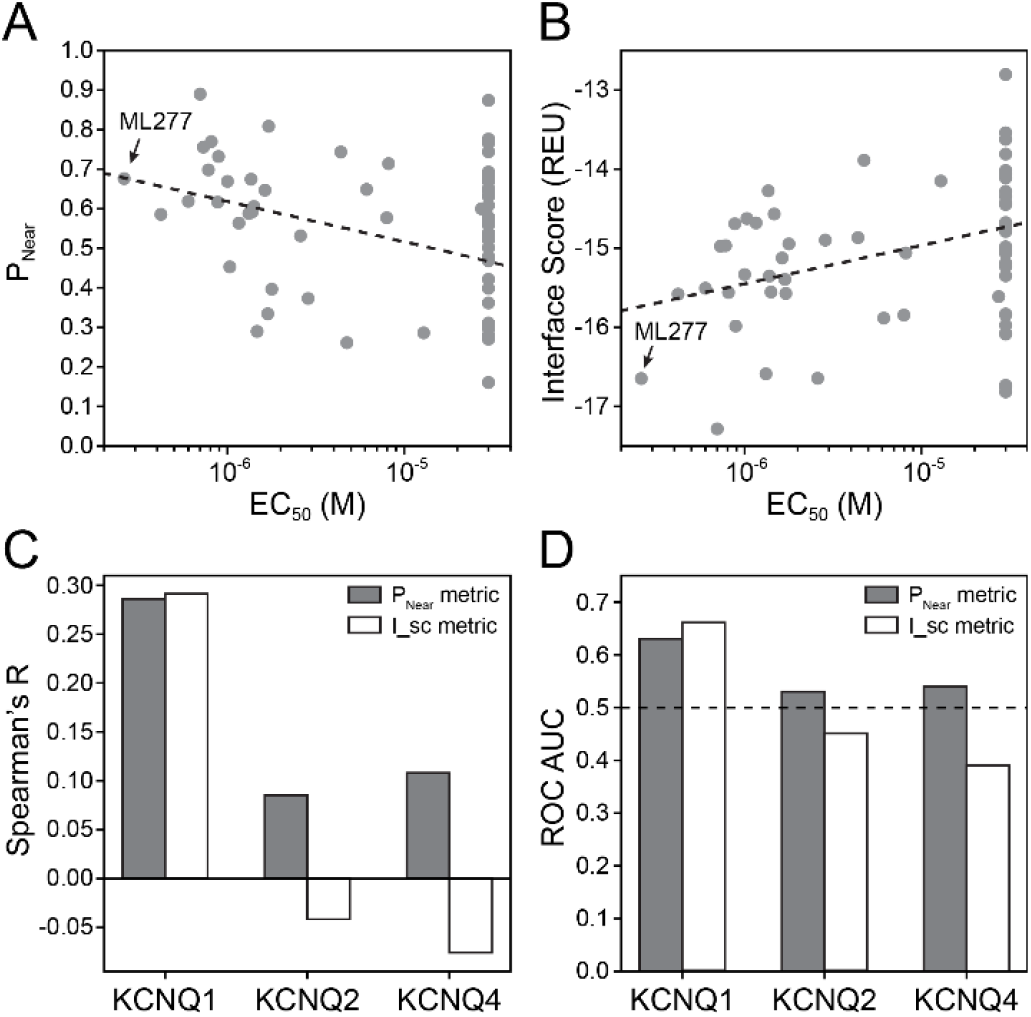
Correlation between docking scores and ligand activity for ML277 and 62 chemical analogs of ML277. (**A**) P_Near_ metric vs. EC_50_ values. (**B**) Rosetta interface score (I_sc) vs. EC_50_ values. (**C**) Spearman’s rank correlation coefficient for the comparison of EC_50_ data with P_Near_ or I_sc scores that were obtained in ligand docking calculations with either KCNQ1, KCNQ2, or KCNQ4 structures, respectively. (**D**) Area under the receiver operating characteristic curve (ROC AUC) for the classification of ligand activity by P_Near_ or I_sc scores that were calculated for either KCNQ1, KCNQ2, or KCNQ4 structures. A ligand activity of EC_50_ = 3 µM was used as binary classification threshold. The AUC of the random baseline classifier is shown as dashed line.

The area under the receiver operating curve (ROC AUC) for discriminating active versus non-active ML277 analogs by their P_Near_ and I_sc values were 63% and 66%, respectively (**Fig 2D**). A significant ranking of ligand EC_50_ data is also observed when the GBVI/WSA scoring function, as implemented in MOE software, was used instead of the RosettaLigand scoring function (Spearman’s R = 0.30, p = 0.017) (**Fig S3C**). By contrast, no discrimination of active versus inactive compounds was observed when ML277 analogs were docked in the same corresponding sites in KCNQ2 and KCNQ4 (**Fig 2C+D**). This is expected since ML277 is not active towards these KCNQ channels and computational docking scores are unrelated to ligand activity in this case. Furthermore, no correlation of ligand EC_50_ with P_Near_ or I_sc was found when ML277 was docked at a slightly different location in KCNQ1 between S2-S3L, S4-S5L, and the loop connecting S4 to S4-S5L (see **Fig S3D**). It was previously suggested that ML277 enters the KCNQ1 channel from the intracellular milieu and that the named site represents the entry point to which ML277 binds first and from which it could reach other regions in KCNQ1 (29). Notably, PIP2 can bind to the same S2-S3L+S4-S5L site. While in our simulations, docking poses are sampled in this region as well, the docking scores were overall lower, and no correlation with ligand SAR data to the extent observed for the S5+S6 cavity site was found. We therefore suggest that the site on S5 and S6 (**Fig 1**) represents the major ML277 binding site that is primarily responsible for ML277 action on KCNQ1. Having identified an ML277 intramembrane binding site in KCNQ1 we next focused on determining the amino acid residues that are important for affinity and selectivity of ML277 towards KCNQ1.

### Energetic analysis of ML277-KCNQ1 residue interactions by MD simulation

To investigate the contributions of individual amino acid residues for ML277 binding, we conducted MD simulations of KCNQ1 closed and open state structures in complex with ML277. The cryo-EM structures of KCNQ1 (15) have the VSD and PD in either activated/closed or activated/open conformations and are referred to as AC and AO state structures. The stoichiometry of the KCNQ1-ML277 complex was 1:1 (i.e., each KCNQ1 subunit was occupied by one ML277 molecule). In addition, we ran MD simulations of the KCNQ1-KCNE1 channel with ML277 bound. KCNQ1-KCNE1 structure models in 1:1 stoichiometry were created by homology modeling. A previously constructed docking model of KCNQ1-KCNE1 (17) and the cryo-EM structure of the KCNQ1-KCNE3 complex (15) were used as templates for homology modeling. ML277 was then docked to the KCNQ1-KCNE1 structure in the same manner as for the KCNQ1 apo structure.

MD simulations of each molecular complex and channel state were run in triplicate for a total length of 1.6 µs. The binding energy of ML277 on KCNQ1 and the per-residue energy contributions were calculated by the MM/PBSA procedure. **Figure 3** displays the MM/PBSA energies along the KCNQ1 amino acid sequence and **Table 1** lists the amino acids with the most negative MM/PBSA energies for all KCNQ1-ML277 complexes analyzed. The MM/PBSA binding energies of all four complexes are similar and not significantly changed by the channel activation state (AC or AO) or the presence or absence of KCNE1. The residues with the largest contribution to the ML277 binding energy are: W248, I268, L271, G272, F275, S276, F279, C331, F332, F335, and A336. As described above, these residues line the ML277 binding site on S5+S6 in KCNQ1.

**Figure 3:**
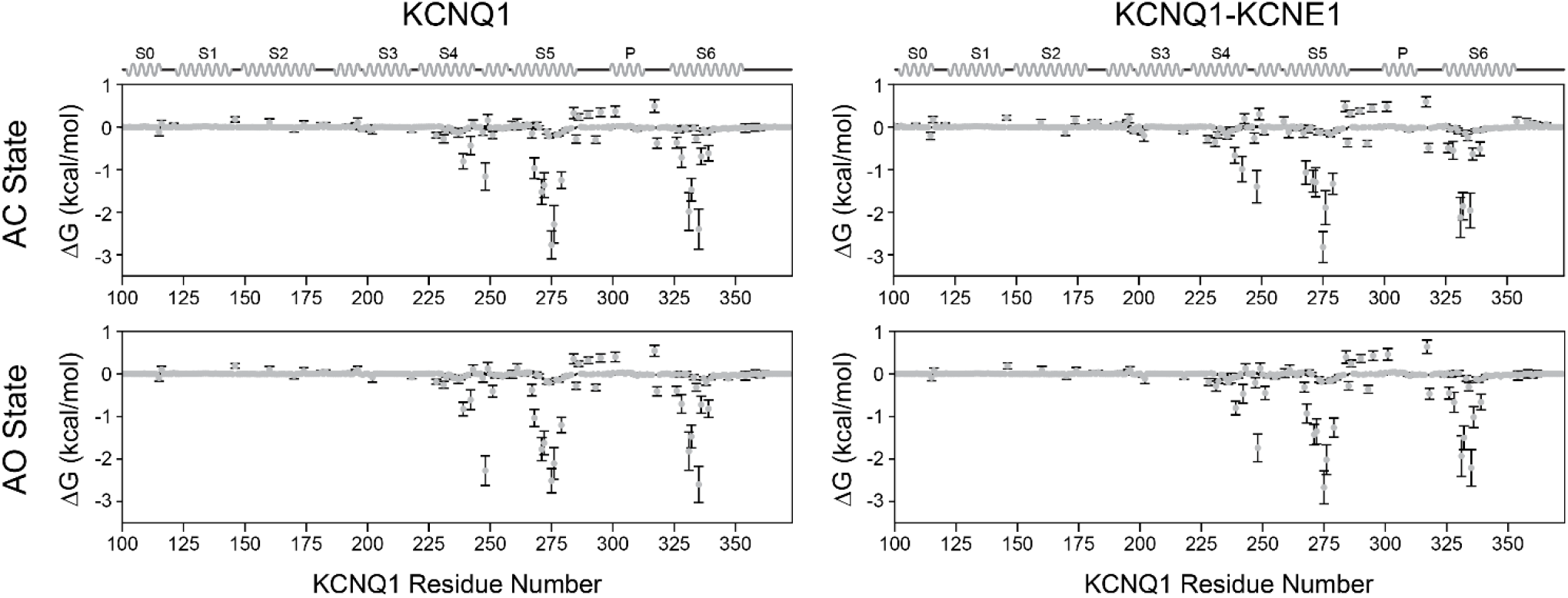
MM/PBSA binding free energy of ML277 per residue of KCNQ1. Values represent the mean ± SD obtained by averaging data over three MD simulation replicas and four KCNQ1 subunits. MM/PBSA energies are shown for the activated-closed (AC) and activated-open (AO) state of KCNQ1 and KCNQ1-KCNE1 channels, respectively. Protein helix regions are depicted and labeled above the plots.

**Table 1:** KCNQ1 residues with largest contributions to ML277 binding energy and their per-residue MM/PBSA energies (in kcal/mol). Values represent mean ± SD and were calculated as average over four KCNQ1 subunits and three MD replicas for each channel state.

|  | KCNQ1 AC | KCNQ1 AO | KCNQ1-KCNE1 AC | KCNQ1-KCNE1 AO |
| --- | --- | --- | --- | --- |
| **Leu239** | -0.81 ± 0.18 | -0.82 ± 0.16 | -0.66 ± 0.18 | -0.80 ± 0.16 |
| **Trp248** | -1.16 ± 0.32 | -2.27 ± 0.35 | -1.40 ± 0.38 | -1.74 ± 0.32 |
| **Ile268** | -1.00 ± 0.24 | -1.04 ± 0.20 | -1.07 ± 0.28 | -0.93 ± 0.22 |
| **Leu271** | -1.52 ± 0.29 | -1.77 ± 0.27 | -1.27 ± 0.26 | -1.43 ± 0.24 |
| **Gly272** | -1.36 ± 0.29 | -1.62 ± 0.28 | -1.30 ± 0.34 | -1.35 ± 0.30 |
| **Phe275** | -2.77 ± 0.33 | -2.51 ± 0.29 | -2.82 ± 0.36 | -2.67 ± 0.39 |
| **Ser276** | -2.29 ± 0.44 | -2.10 ± 0.37 | -1.89 ± 0.40 | -2.02 ± 0.35 |
| **Phe279** | -1.25 ± 0.19 | -1.20 ± 0.18 | -1.33 ± 0.25 | -1.26 ± 0.22 |
| **Cys331** | -1.99 ± 0.44 | -1.82 ± 0.45 | -2.12 ± 0.47 | -1.93 ± 0.48 |
| **Phe332** | -1.47 ± 0.27 | -1.47 ± 0.27 | -1.86 ± 0.32 | -1.50 ± 0.28 |
| **Phe335** | -2.40 ± 0.47 | -2.60 ± 0.43 | -1.96 ± 0.41 | -2.21 ± 0.43 |
| **Ala336** | -0.69 ± 0.19 | -0.72 ± 0.20 | -0.63 ± 0.14 | -1.02 ± 0.25 |

The finding that residues on S5 and S6 contribute strongly to the ML277 binding energy is in line with previous mutagenesis and electrophysiology studies (29), which showed that mutations of residues F275, F332, and F335 in the PD to alanine decrease the effect of ML277 on KCNQ1. Furthermore, our simulations identified additional residues that could be critical for binding, such as G272, S276, and F279.

We superimposed the structure of KCNQ1 with those of KCNQ2 (33) and KCNQ4 (34) and this revealed several amino acid differences in the region predicted to be the ML277 binding site (see **Fig S4**). Notably, G272 in KCNQ1 is substituted by cysteine and valine in KCNQ2 and KCNQ4, respectively. These larger amino acids are expected to narrow the ML277 binding site and potentially clash with ML277, preventing ligand binding. Another KCNQ1 residue, S276, is replaced by alanine in KCNQ2 and KCNQ4, an amino acid that fails to form a hydrogen bond to ML277 and therefore is expected to further attenuate ML277 binding. Furthermore, several phenylalanine residues in KCNQ1 are replaced by leucines in KCNQ2 and KCNQ4, which is also expected to decrease the interaction with ML277. Specifically, F275, F279, and F335 in KCNQ1 are leucines in KCNQ2, and F279 and F335 are leucines in KCNQ4. Moreover, C331 and A336 are substituted by small or more polar amino acids in KCNQ2 and KCNQ4: C331 in KCNQ1 is threonine in KCNQ2 and glycine in KCNQ4, while A336 is glycine in both KCNQ2 and KCNQ4. Overall, the amino acids in KCNQ2 and KCNQ4 appear to provide less favorable binding sites for ML277, which could explain why these channels are insensitive to ML277.

### Functional analysis of KCNQ channel subtype residue differences on ML277 effects

To solidify our observations from the computational simulations and investigate the molecular basis for selectivity of ML277 for KCNQ1 over KCNQ2 and KCNQ4, we carried out site-directed mutagenesis experiments on KCNQ1 coupled to electrophysiology of the designed KCNQ1 mutants. We designed mutations that swap amino acids in KCNQ1 to the corresponding amino acids in KCNQ2 or KCNQ4. We generated the G272V, F275L, S276A, F279L, C331G, A336G mutants that are located in the predicted ML277 binding site. These channels were transiently expressed in CHO cells stably expressing KCNE1 (CHO-E1 cells) and their consequences were determined by measuring ML277-induced effects on peak and tail current density, voltage-dependence of activation V_1/2_ and deactivation rate, three channel properties previously described as targets of ML277 modulation (28–31).

We first determined the functional properties of individual KCNQ1 mutants under control conditions without ML277. The six mutant channels exhibited distinct peak and tail current density, voltage-dependence of activation and deactivation rates compared to the wild type (WT) channel. F279L displayed larger JNJ303-sensitive currents than WT, while the other five mutant channels exhibited smaller current densities. The S276V and F279L mutant channels showed depolarized V_1/2_, while the V_1/2_ calculated for F275V was hyperpolarized. No change was observed for G272V, while the tail currents measured for C331G and A336G were too small for reliable determination of V_1/2_. Deactivation kinetics were slower for G272V, F275V and S276V, faster for F279L and C331G, and unchanged for A336G. These results are summarized in supporting **Table S1. Figure S5** shows the average whole-cell currents recorded before and after JNJ303 addition, and the JNJ-sensitive currents.

The effects of ML277 on current density, V_1/2_, and channel deactivation rate were assessed by comparing these properties measured for each mutant channel in the absence (control) and presence of ML277. **Figure 4A** shows the average peak and tail whole-cell, JNJ303-sensitive current density, recorded from CHO-E1 cells expressing WT or mutant KCNQ1 channels exposed to vehicle (control, black line) or ML277 (blue line) and normalized to membrane capacitance. The vertical scale for each channel trace was modified to better illustrate the effect of ML277. Statically significant (p ≤ 0.02) increases in current density due to ML277 exposure were measured for WT (peak and tail) and F275V (tail) channels. No significant effects on current density were observed for the G272V, F279L, C331G and A336G mutant channels. Interestingly, the mutant S276V channel exhibited a significant ML277-dependent decrease in tail current (**Fig 4A**). Significant effects of ML277 on voltage-dependence of activation V_1/2_ were seen only for the F279L mutant channel (-8 mV, **Fig 4C**). Under the conditions used in this study, the WT channel exhibited a smaller, less significant hyperpolarizing shift in V_1/2_ (∼-2 mV, p = 0.038). Exposure to ML277 slowed channel deactivation in four out of six mutant channels (G272V, S276A, F279L and C331G), while having the opposite effect on F275V. No effects were observed on the deactivation of A336G.

**Figure 4:**
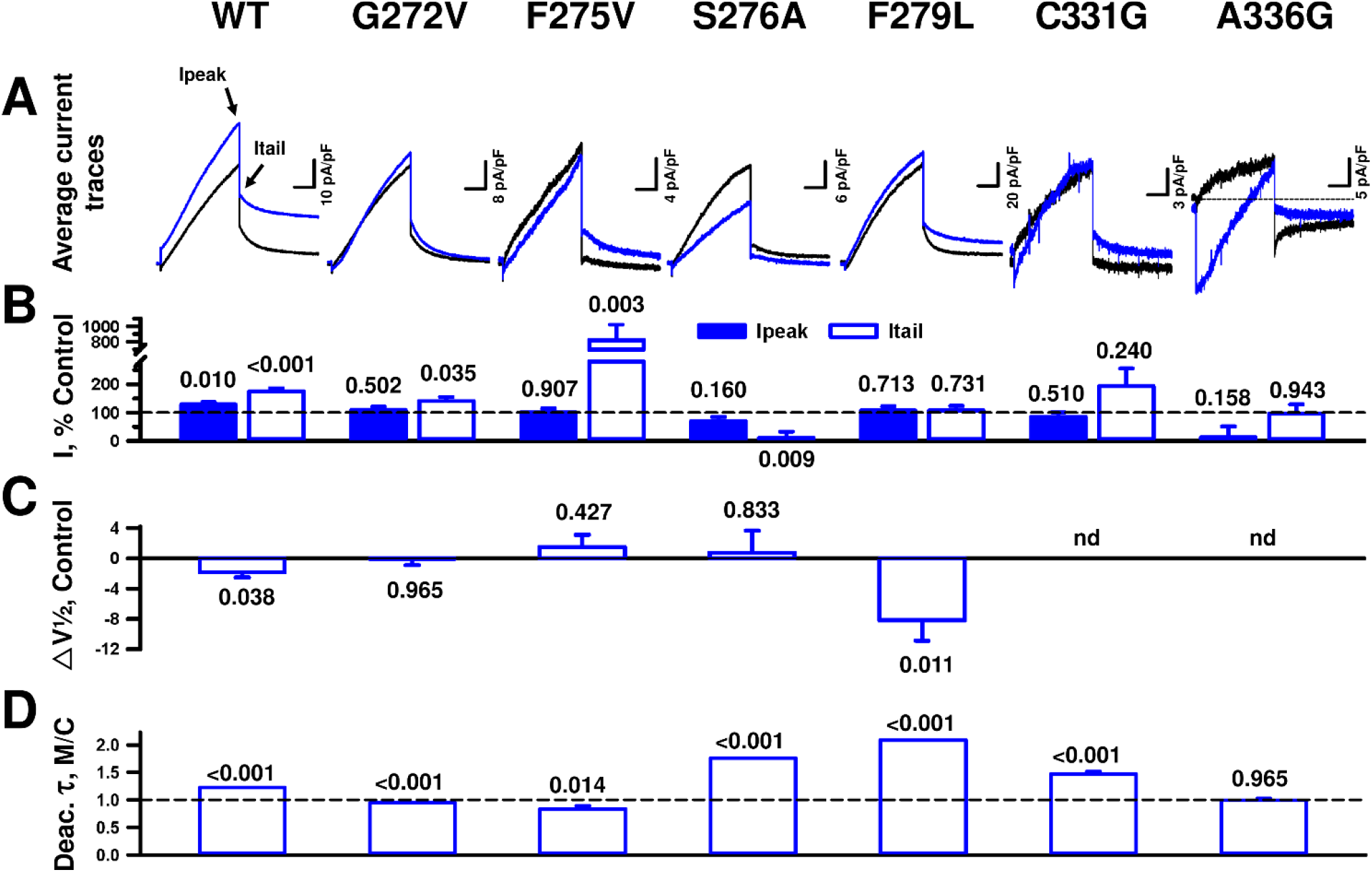
Impact of mutations in amino acid residues located in the predicted ML277 binding site on ML277-induced effect. **A** Average whole-cell, JNJ303-sensitive currents recorded at +60 mV from CHO-E1 cells transiently expressing WT or mutant KCNQ1 channels exposed to vehicle (black) or ML277 (blue). Arrows indicate the time points where peak current (1990 ms after start of first pulse) and tail current (10 ms after start of -30 mV pulse) were measured. Horizontal scale bar represents 500 ms, and the dashed line indicates zero current (A336G). **B**. ML277-dependent change in peak current density (empty bars) or tail current density (solid bars) relative to control (vehicle) cells recorded in parallel. **C**. ML277-induced change in voltage-dependence of activation V½ relative to control cells recorded in parallel. **D**. Change in channel deactivation time constant induced by ML277 expressed as the ratio of values determined in the presence (M) or absence (C) of the compound. Statistical analysis was performed by *t*-test, and P values are shown in the figure. Number of cells analyzed: WT: control = 96-156, +ML277 = 121-156; G272V: control = 37-57, +ML277 = 49-63; F275L: control = 10-47, +ML277 = 13-48; S276A: control = 35-69, +ML277 = 20-62; F279L: control = 24-23, +ML277 = 20-26; C331G: control = 42, +ML277 = 42; A336G: control = 13, +ML277 = 12.

**Table S2** summarizes the results shown in **Figure 4** while **Figure S6** depicts the average whole-cell current recorded for all channels in the presence of ML277, before and after JNJ303 addition, and the JNJ303-sensitive currents. The F279L mutant exhibited the most interesting effects. This phenylalanine to leucine substitution augments the ML277-induced effects: larger hyperpolarizing shift in V_1/2_ and slower deactivation than observed for the WT channel. The apparent absence of current density increase due to ML277 may be explained by the observation that this mutant channel appears more resistant to JNJ303 block (see **Fig S6**).

In summary, most of the mutations introduced prevented and even reversed the expected effect of ML277 on I_Ks_ channel activity. ML277-dependent increase in current density and hyperpolarizing shift in voltage-dependence of activation V_1/2_ were most susceptible to mutations. G272V, F275V, S276A, F279L, C331G, and A336G prevented the ML277-indiuced increase in peak current, and G272V, F275V, S276A prevented the hyperpolarizing shift of V_1/2_, while tail currents of C331G and A336G were too small for reliable V_1/2_ measurement. ML277-evoked changes in deactivation were the most resistant to disruption by the mutations.

### MD network analysis reveals that ML277 enhances allosteric signaling between the KCNQ1 VSD and PD

We next investigated how ML277 modulates KCNQ1 molecular function by interacting with the identified binding site. Previous studies have provided evidence that ML277 activates KCNQ1 by promoting VSD-PD coupling when the VSD is in the activated state to increase the current of the AO state (30). In VSD-PD coupling, movements of S4 are transmitted to the PD through a chain of residue-residue interactions that constitute allosteric pathways. Allosteric signaling then triggers conformational changes in the PD that lead to channel opening. To understand how ML277 influences these allosteric pathways we employed network analysis of our MD simulations. This approach was previously applied to reveal the specifics of calmodulin modulation of KCNQ1 allostery (13) and was recently extended to include lipids and other small-molecule cofactors in the analysis (45).

The concept of network analysis is outlined in **Figure 5A**. MD trajectories of KCNQ1 or KCNQ1-KCNE1 with ML277 were converted into networks by assigning a node to every protein residue. Lipids and ML277 were included as additional interactors in the network; three nodes were assigned to every POPC residue and three nodes were assigned to every ML277 molecule. The node assignment of atoms in POPC and ML277 is shown in **Figure S7**. The connections (edges) between network nodes were assigned weights that capture the strength of residue interactions in KCNQ1 or between KCNQ1 and cofactors. The weights were defined based on the spatial proximity of residues and the degree of correlated residue motions. In the final network, pathways connecting the VSD gating charges (i.e., the source residues) to the pore-gate residue S349 (i.e., the sink residue) were extracted and the amount of information flow along those pathways was calculated taking into account the weights across all pathway edges. Thus, residues and pathways that are important for VSD-PD coupling are highlighted by increased information flow. Meanwhile, uninteresting protein regions are characterized by low information flow because pathways move back and forth across edges such that information transmission is unproductive.

**Figure 5:**
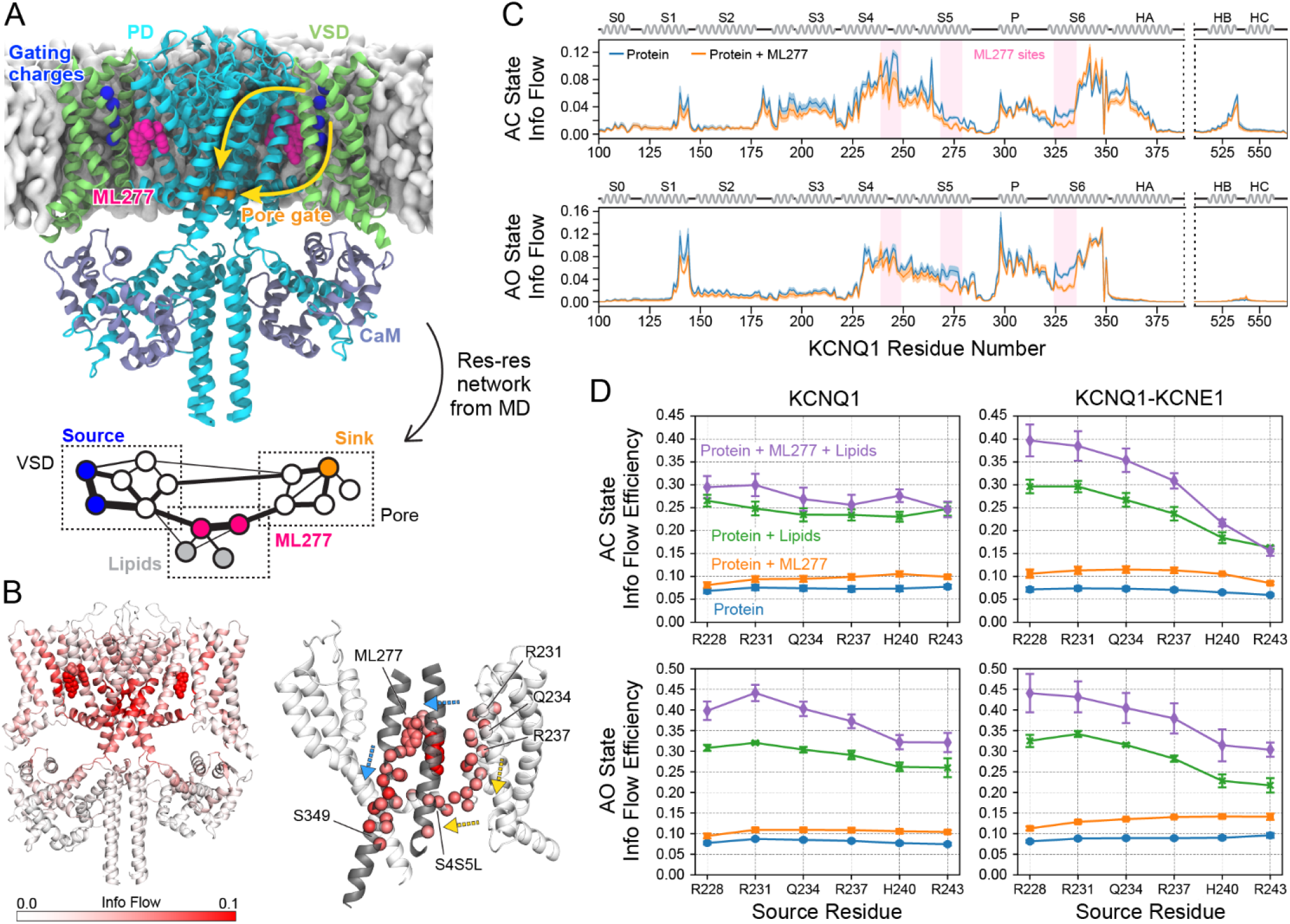
MD network analysis demonstrates that ML277 modulates VSD-PD allosteric pathways in KCNQ1. (**A**) Molecular components and schematic network representation of MD system. Two complete KCNQ1-CaM subunits are shown. The VSDs, CaMs, and HA and HB helices in the front and back are omitted. The protein, ML277, and lipids are transformed into a network in which each residue and cofactor molecule represents a node and each edge weight corresponds to the interaction strength between residues. Network analysis is then applied to trace allosteric pathways and determine information flow efficiency from the VSD gating charges (i.e., source residues) to the PD gate (i.e., sink residues). (**B**) Left: Information flow through all pathways connecting the VSD to the PD gate is projected onto the KCNQ1 AC structure. Dark red corresponds to high information flow (capped at 0.1). Right: Residues with high information flow along the pathways between the S4 gating charges and the PD gate are depicted as spheres. Only the intramembrane part of one KCNQ1 subunit (white) and S5 and S6 from a neighboring subunit (grey) are shown. The direction of two major pathways, one involving ML277 (blue arrows) and another one involving S4-S5L (yellow arrows) are shown. (**C**) Information flow through KCNQ1 AC and AO structures with and without ML277 bound along the residue sequence. Changes in information flow in the presence of ML277 are observed mostly for residues in the ML277 binding regions indicated by light pink shadings. (**D**) Information flow efficiency for pathways connecting the source residues listed on the x-axis to the sink residue S349. Information flow efficiency is shown for networks containing KCNQ1 protein alone or together with ML277 and/or lipids. Information flow efficiency is displayed for AC and AO state structures of KCNQ1 and KCNQ1-KCNE1. Information flow of protein + lipids was measured from simulations that were conducted without ML277 under the same conditions.

**Figure 5B** visualizes the spatial pattern of allosteric pathways in KCNQ1 by projecting the calculated information flow onto the 3D structure of KCNQ1+ML277. **Figure 5C** displays the information flow for every KCNQ1 residue in the AC and AO state structure with or without ML277. As expected, the highest information flow residues in both conformations are observed along two pathways connecting the VSD gating charges to S349. One pathway involves S4, S4-S5L, and the lower half of S6 and is considered the canonical coupling pathway. Another major pathway involves the interface between S4 and S5 from two neighboring subunits and the S6 helix. This alternative coupling pathway has only recently been identified in K_V_ channels (46). The two pathways are highlighted by yellow and blue arrows in **Figure 5B**. They have been studied in domain-swapped K_V_ channels and also identified in KCNQ1 before (11–13).

Looking at ML277, high information flow through the ligand is recognized, as clearly seen by the dark red color of ML277 in **Figure 5B**. It is further recognized that ML277 is part of a chain of high information flow residues. The chain starts with the first gating charge residues on S4 (R231, Q234, R237), runs through ML277, and propagates down S6 to reach the pore gate residue S349. Notably, the residues along this ML277-mediated pathway exhibit high information flow values of >0.05, highlighting the importance of this pathway for allosteric signaling. We next investigated whether the MD simulations could reveal a coupling-enhancing effect of ML277. Simulations in ML277-bound and ML277-free states yielded similar information flow profiles for KCNQ1, for both the AC and the AO state structure (see **Fig 5C**). Surprisingly, the presence of ML277 leads to a decrease of information flow for residues in regions that interact with ML277. This is due to the fact that these amino acid residues are bypassed by ML277 in the network and that pathways are rerouted through ML277. Hence, by looking only at KCNQ1, a signaling enhancing effect is not apparent. We thus turned to the metric of information flow efficiency closeness which is a measure of signaling efficiency in the whole network, including lipids and cofactors. **Figure 5D** shows signaling efficiency for pathways connecting each of the six S4 gating charge residues to S349. Indeed, a clear increase of signaling efficiency is observed when ML277 is present. This effect is even stronger when lipids are included in the network. The highest efficiency is found when both lipids and ML277 are part of the network. This effect is consistently observed under all simulation conditions independent of the KCNQ1 conformation state (AC or AO) or the presence or absence of KCNE1. This finding suggests that KCNQ1 exploits ML277 and surrounding lipids for promoting allosteric signaling between the VSD and PD. Tightly bound interactors that snuggle into surface cavities between the VSD and PD may provide a mechanism to increase allosteric efficiency in KCNQ1. If this is the case, then we expect that chemical analogs of ML277, which fail to activate KCNQ1 *in vitro*, will also fail to increase information flow efficiency to the same extent as seen for ML277 because they bind too weakly or lack favorable interactions with KCNQ1. To test this hypothesis, we performed additional MD simulations of KCNQ1 AC and AO state structures with two inactive analogs of ML277 – the *S*-enantiomer of ML277 and an analog in which the terminal methylphenyl group is changed to a chemically different chlorobenzyl group. The chemical structures of these analogs are displayed in **Fig 6A**. As expected, the inactive *S*-enantiomer of ML277 does not increase the information flow efficiency and the drug bound channel behaves similar as the apo state (**Fig 6B**). The second analog induces only a moderate increase of the information flow efficiency to an extent that is much weaker than for ML277, especially in the AO conformation (**Fig 6C**). These observations are consistent with the much lower activity of these analogs of EC_50_ >30 µM compared to EC_50_ = 0.26 µM for ML277 (27). In summary, we conclude that ML277 acts as an allosteric activator of KCNQ1 by serving as a hub in an allosteric pathway that connects S4 with S5 and S6 to increase the efficiency of VSD-PD coupling.

**Figure 6:**
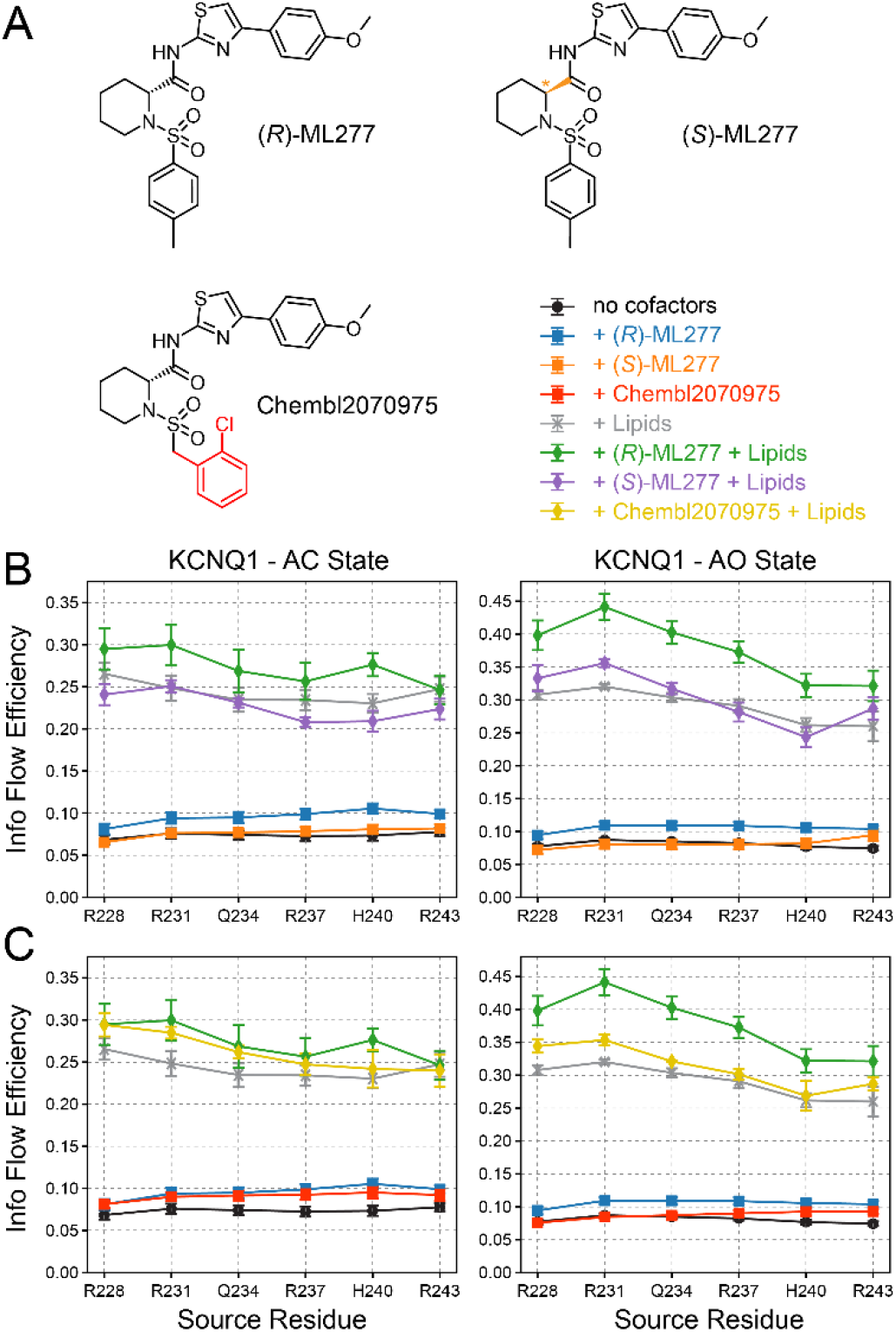
Inactive analogs of ML277 fail to increase information flow efficiency between KCNQ1 VSD and PD. (**A**) Chemical structures of the *R*-and *S*-enantiomers of ML277 and of molecule CHEMBL2070975. (**B**) Information flow efficiency for pathways connecting the source residues listed on the x-axis to the sink residue S349 in MD simulations of KCNQ1 alone or with either (*R*)-ML277 or (*S*)-ML277. Information flow efficiency is displayed for the AC and AO state structures of KCNQ1. (**C**) Information flow efficiency for pathways between the source residues listed on the x-axis to the sink residue S349 in MD simulations of KCNQ1 alone or with either (*R*)-ML277 or CHEMBL2070975.

## Discussion

In this study, we identified an intramembrane binding site for the small-molecule KCNQ1 activator ML277. Using computational docking and MD simulations, we investigated the molecular mechanism of ML277 action and corroborated the deduced ML277 binding model by complementary computational and experimental approaches. First, computationally predicted binding energies for ML277 and 62 chemical analogs of ML277 could be successfully used to rank the structure-activity relationships for this set of compounds (27). This was not possible when ML277 was docked at a different region in KCNQ1 or at the same region in KCNQ2 or KCNQ4 channels. These results suggest that our binding model is able to capture the specifics of KCNQ1-ML277 interaction and can be helpful to interpret the impact of chemical and structural modifications on drug activity. Second, KCNQ1 residues involved in ML277 binding were pinpointed by MM/PBSA energy analysis and experimentally tested by site-directed mutagenesis. Specific amino acid mutations either reduced the ML277 effect or profoundly changed the KCNQ1 response to ML277. This is in line with the notion that these amino acid residues take part in binding of ML277 or are involved in regulation of allosteric coupling by ML277. Third, MD simulations indicated that that ML277 modulates allosteric networks in KCNQ1 and increases the efficiency of allosteric signaling between the VSD and the PD. This result complements previous functional studies (11, 30, 31) showing that ML277 enhances VSD-PD coupling to increase KCNQ1 channel conductivity, specifically when the channel adopts the AO state. The MD simulations provided an atomic-level picture of the coupling process and a means to evaluate drug action. This level of information is not available from the inspection of static structures. Notably, analogs of ML277, which have no detectable activity on KCNQ1 (e.g., the (*S*)-enantiomer of ML277), were found to lack favorable interactions with KCNQ1 and failed to significantly increase signaling efficiency in MD simulations. Thus, the MD network approach proved to be a powerful tool to obtain functional insights into drug action from ion channel structures.

The ML277 binding model is consistent with the results of structure-function studies on ML277 (11, 29). Work by Xu et al. (29) and Hou et al. (11) found 18 mutations in KCNQ1 that abolished the ML277 effect. This could be explained by mutations disrupting ML277 binding or because they impaired VSD-PD coupling. **Figure 7** shows the location of those 18 mutations and of the additional six mutations identified in this study, mapped onto the KCNQ1 sequence. There are 21 critical mutation sites; 15 of these sites correspond to residues that line the ML277 binding pocket in the structural model (i.e., these residues have a heavy-atom distance to ML277 smaller than 4.5 Å). The other six mutation sites are either directly adjacent to a ML277 binding residue or are in regions known to be critical for VSD-PD coupling such as the S4-S5 linker. Altogether these data provide strong evidence that ML277 binds at the interface between S4, S4-S5L, S5, and S6, as we have corroborated using our structural model for ML277 binding.

**Figure 7:**
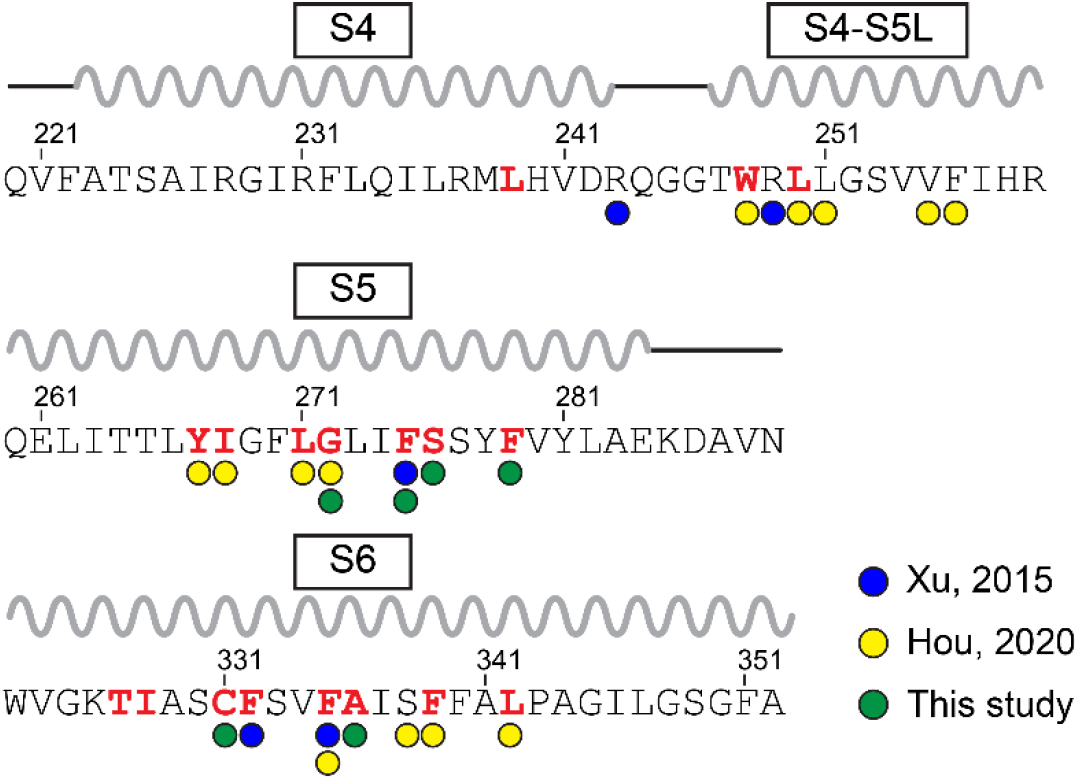
Location of predicted ML277 binding site residues and of mutations that reduce the ML277 effect on KCNQ1. ML277 binding site residues are highlighted in bold red font. The positions of high-impact mutations, which were reported in the literature (11, 29) or found in this study, are shown with colored circles below the amino acid sequence.

ML277 has a favorable selectivity profile and is >100-fold more active on KCNQ1 than on the related KCNQ2 and KCNQ4 channels (27). It is interesting that amongst the ML277 binding residues displayed in **Fig 1D**, nine residues have a different identity in KCNQ2 and eight are different amino acids in KCNQ4 (see **Fig S4**). Overall, these divergent amino acids seem to provide less favorable interactions for ML277. This offers a plausible explanation for why ML277 is less active on KCNQ2 and KCNQ4. G272 is changed to cysteine and valine in KCNQ2 and KCNQ4, respectively. These larger amino acids are expected to narrow the ML277 binding pocket, which could prevent ligand binding. Another KCNQ1 residue, S276, is substituted by alanine in KCNQ2 and KCNQ4. Alanine fails to form side chain hydrogen bonds, which could further reduce ML277 binding. Furthermore, F275 is changed to leucine in KCNQ2, and F279 and F335 correspond to leucine in both KCNQ2 and KCNQ4. These amino acid substitutions reduce the hydrophobic surface area and could prevent stacking interactions with aromatic rings in ML277. Moreover, C331 is changed to threonine and glycine in KCNQ2 and KCNQ4, respectively, and A336 is swapped with glycine in both KCNQ2 and KCNQ4. This also reduces the hydrophobic surface area and could impair ML277 binding. These structural interpretations are consistent with the functional results obtained with the KCNQ1 mutants tested in this work. Mutating G272, F275, S276, F279, C331, and A336 to their corresponding amino acids in KCNQ2 or KCNQ4, either reduced or profoundly changed the ML277 effect on KCNQ1. The reduced or absent effect by ML277 is likely not due to an excess of KCNE1 expression relative to KCNQ1 expression previously shown by Yu et al (28). Co-transfection of KCNQ1 with KCNE1 into the CHO-E1 cell line generated larger currents than when KCNQ1 was transfected alone.

Our results for the F279L mutant suggest an increased sensitivity to ML277 binding, not the expected reduced or changed response. This augmented response may be due to increased channel affinity for ML277 or other mechanisms. For example, previous studies by Nakajo and Kubo (18) indicated that F279 interacts with F232 when KCNE1 is expressed to destabilize the open state. Thus, either the amino acid substitution or ML277 binding may disrupt the interaction between these two residues, generating the stronger response observed. A more stable open state in F279L channel may explain the decreased block by JNJ303. JNJ303 is an adamantane derivative suggested to bind to the I_Ks_ channel close state (47). We also observed a decreased block by JNJ303 on the mutant A336G. Previous studies showed that mutating this residue increased the IC_50_ for JNJ303 ∼4-fold (47).

Previous studies demonstrated that the activity of ML277 is dependent on the presence of KCNE1 (28–31). The ML277 effect was reduced with increasing expression levels of KCNE1. Saturating KCNE1 levels led to little (29) or no (28) effect of ML277 on whole-cell currents and abolished the ML277-induced slowing of channel deactivation. On the other hand, at the single channel level, KCNQ1-KCNE1 channels were still potentiated by ML277 (31). Channel openings were more frequent and had larger amplitudes in presence of ML277, but were more flickery compared to openings of single KCNQ1 channels with ML277 (31). Our simulation data provide some hints on how KCNE1 could change ML277 action. We observed that ML277 was able to bind to the same site on KCNQ1 under KCNE1-free and KCNE1-bound conditions and that there was no steric overlap of the ML277 and KCNE1 binding sites. Only a single side chain contact between ML277 and KCNE1-F53 was observed in MD (**Fig S8**), but this did not disturb ML277 binding. Hence, it is unlikely that KCNE1 and ML277 compete for binding. Looking at the MD network analysis and comparing the information flow through the KCNQ1 and KCNQ1-KCNE1 channel structures we found that KCNE1 profoundly changes allosteric signaling in KCNQ1, especially in the AO state structure (**Fig S9**). Information flow is reduced mostly in areas of the central pore domain, but is increased in regions of the VSD and at the S6-HA connector helix (**Fig S9**). Information flow through ML277 itself is reduced as well. This suggests that the coupling pathways linking S4 to S5+S6, which are promoted by ML277 in the KCNE1-free structure, become less important for signaling when KCNE1 is bound. This would render ML277 less active in promoting VSD-PD coupling and could explain the loss ML277 activity for the KCNQ1-KCNE1 complex. However, more investigations will be needed to fully resolve this issue. Network analysis is expected to provide an important tool to guide the design of functional experiments that will test the impact of KCNE1 and other auxiliary proteins on signaling in KCNQ1.

It is interesting that ML277 and another KCNQ1 activator, R-L3, share overlapping binding sites and have similar effects on KCNQ1 (20, 48). R-L3 increases current amplitude, shifts the activation voltage to more hyperpolarized potentials, and slows channel activation and deactivation. Co-expression of KCNQ1 with KCNE1 reduces the R-L3 effect, and at saturating KCNE1 levels no R-L3 effect is observed (20), similar to what was seen for ML277. Using scanning mutagenesis, the putative binding site for R-L3 was localized to residues in S5 and S6. Mutations at Y267, I268, L271, G272, F335, and I337 caused the most significant decrease in drug sensitivity (48). Recently, a molecular binding model for R-L3 was developed and additional interactions with the VSD were confirmed (49). Interestingly, the R-L3 binding residues are part of the putative ML277 binding site. This suggests that R-L3 and ML277 have related mechanisms of action, even though they have different chemical structures. Moreover, it could become possible to develop additional small-molecule modulators that target the same area in KCNQ1. In fact, the observation that small ligands and auxiliary lipids and proteins recognize common regions in KCNQ1 and other ion channels that are important for electromechanical coupling, underscores the possibility to harness these sites for regulating channel activity. Recently, Liu et al. (50) identified a synthetic compound (CP1) that can substitute for PIP2 and mediates VSD-PD coupling by recognizing a binding site on KCNQ1 that overlaps with that of PIP2. CP1 could rescue channel rundown induced by PIP2 depletion and shorten the action potential in guinea pig ventricular cardiomyocytes (50). These results highlight the potential of this strategy for KCNQ1 modulator design. Furthermore, several examples of intramembrane small-molecule binding sites have been found in ion channel structures lately (51). This motivates the development of new concepts and considerations for drug design efforts targeting these sites.

## Methods

### Structure preparation

The cryo-EM-determined structures of KCNQ1 in the AC state (PDB: 6UZZ, 6V00) and AO state (PDB: 6V01) (15), respectively, were used as receptor models for ligand docking. Missing residues between helix S3 and S4 in KCNQ1 (G219-F222 for PDB 6UZZ and 6V00, G219-T224 for PDB 6V01) were added with RosettaRemodel (52). The protein structure was minimized with Rosetta (version 3.12) (40) using the RosettaMembrane energy function (53, 54) and an additional score term reflecting the fit to the experimental electron density map (EMD: 20965, 20966, 20967). During minimization and relaxation into electron density, C4 symmetry was enforced with the help of Rosetta symmetry definition files (55).

### Ligand docking

Ligand docking was performed with RosettaLigand (39, 56) through RosettaScripts (57) as described previously (58). Rosetta-formatted ligand parameter files and ligand conformers were generated using the BCL software (59) as detailed in Moretti et al. (60). To find low-energy binding modes of ML277 in KCNQ1, ligand docking was carried out in two steps: (1) ML277 was placed at multiple starting positions in a side pocket in KCNQ1 that is surrounded by S2-S3L and S4-S5L on the intracellular side and helices S4, S5, and S6 on the lateral sides. Previously, Xu et al. (29) localized the ML277 binding region to this side pocket based on the observation that mutations of residues in S2-S3L, S4-S5L, and S4-S6 reduce the ML277 effect. A total of 20,000 ML277 docking poses were generated in this step. (2) Representative ML277 docking poses were identified by hierarchical clustering. A root mean-square distance deviation (RMSD) cutoff of 4 Å between two ligand poses was used to define a cluster. The models with the best interface scores from the ten largest clusters served as starting point for one more round of local ligand docking. ML277 was redocked 3,000 times into KCNQ1 in this refinement step. Representative binding models of ML277 were identified by interface score and P_near_ metric (61), which evaluate the binding energy of a docked ligand and the energy gap between the best-scoring pose and other similar docking poses, respectively. Ligand docking was repeated for a series of 62 chemical analogs of ML277, and docking scores were compared to EC_50_ data available for these compounds (27). Prior to docking, each analog was aligned to the ML277 binding pose using BCL::MolAlign (62) and afterwards re-docked into KCNQ1 3,000 times using RosettaLigand (39, 56).

Control ligand docking calculations were performed with Molecular Operating Environment (MOE) 2020.09 (Chemical Computing Group ULC, Montreal, QC, Canada). For ML277 and each of its 62 analogs, ligand conformers were generated with the bond-rotation method in MOE and ligand molecules were placed 5,000 times with the Triangle Matcher method. The top 100 poses after scoring with the London dG scoring function were minimized with flexibility on the receptor and ligand sides enabled. Final poses were ranked using the GBVI/WSA dG scoring function (63).

### MD simulations

MD simulations of KCNQ1 or KCNQ1-KCNE1 channels with Calmodulin (CaM) and with or without ML277 were performed in POPC (palmitoyloleoyl-phosphatidylcholine) membrane bilayers at 303 K using AMBER20. KCNQ1 models were embedded in a bilayer of ∼380 POPC molecules using the membrane builder tool of the CHARMM-GUI website (64). A TIP3P water layer containing 100 mM neutralizing KCl and extending 20 Å from the closest protein atom along the Z axis was added on either side of the membrane. In addition, three K^+^ ions were placed in the channel selectivity filter at positions S0, S2 and S4, while two water molecules were placed at positions S1 and S3. The ff14SB force field for proteins (65) and the Lipid17 force field for POPC (66) were used for all simulations. The geometry of the ML277 molecule was optimized with Gaussian 09 (Gaussian, Inc., Wallingford CT) on the B3LYP/6-31G* level of theory, and assignment of Amber atom types and calculation of RESP charges was done with Antechamber (67). SHAKE (68) bond length constraints were applied to all bonds involving hydrogens. Nonbonded interactions were evaluated with a 10 Å cutoff, and electrostatic interactions were calculated by the particle-mesh Ewald method (69).

Each MD system was first minimized for 10,000 steps using steepest descent followed by 5,000 steps of conjugate gradient minimization. With protein, ligand, and ions restrained to their initial coordinates, the lipid and water were heated to 50 K over 1000 steps with a step size of 1 fs in the NVT ensemble using Langevin dynamics with a rapid collision frequency of 10,000 ps^-1^. The system was then heated to 100 K over 50,000 steps with a collision frequency of 1000 ps^-1^ and finally to 303 K over 200,000 steps and a collision frequency of 100 ps^-1^. After changing to the NPT ensemble, restraints on ions were gradually removed over 500 ps and the system was equilibrated for another 5 ns at 303 K with weak positional restraints (with a force constant of 1 kcal mol^-1^ Å^-2^) applied to protein Cα atoms and ligand heavy atoms. The restraints were then gradually removed over 10 ns, and production MD was conducted for 550 ns using a step size of 2 fs, constant pressure periodic boundary conditions, anisotropic pressure scaling, and Langevin dynamics. Two to four independent simulations were carried out for each state (AC or AO) and each complex (KCNQ1-CaM or KCNQ1-KCNE1-CaM, with or without ML277).

### MD network analysis

Residue interaction networks including protein, lipids, and ML277 were calculated and allosteric pathways were identified using the Allopath tool (45). A node in the network was assigned to every protein residue, three nodes were assigned to every POPC residue – one centered at the PC head group, one centered at the oleoyl chain, and one centered at the palmitoyl chain – and three nodes were assigned to every ML277 molecule. The grouping of cofactor (i.e., lipid and ligand) atoms into domains, which provided the basis for their node assignments, is shown in **Figure S7**. The network was obtained from the elementwise product of a contact map *C* and mutual information map *M*. Calculation of *C* and *M* was carried out as detailed in (45) and by using default parameters for the Allopath tool. MD snapshots were taken at very 0.5 ns from the last 500 ns of the simulation. The obtained networks were analyzed by current flow betweenness analysis to identify allosteric pathways between the VSD and PD and by current flow closeness analysis to estimate the efficiency of information flow along allosteric pathways. The gating charge residues in the VSD (R228, R231, Q234, R237, R240, and R243) were defined as source residues and S349 in the PD was designated the sink residue.

### MM/PBSA calculations

The binding energy (ΔG_binding_) between KCNQ1 and ML277 and the per-residue contributions to ΔG_binding_ were computed with the help of the *MMPBSA*.*py* program (70). A total of ∼1150 MD frames sampled at 450 ps intervals from the last 520 ns of each MD replica were processed to compute the molecular mechanics potential energies and solvation free energies in the MM/PBSA procedure (71). The solvation free energy contribution to ΔG_binding_ was calculated using a continuum Poisson-Boltzmann (PB) model for channel proteins as described in (72). The linear PB equation was applied in calculating the electrostatic part of the solvation energy and the non-polar solvation energy was modeled as a combination of a cavity and dispersion term. Values of ionic radii and charges optimized for PBSA calculations with AMBER atom types (73) were employed and the ionic strength of water was set to 100 mM. PBSA calculations were carried out using a finite-difference grid with a spacing of 0.5 Å and a fill ration of 1.5 (i.e., the longest dimension of the grid over the longest dimension of the solute). The geometric multigrid solver with a convergence threshold of 0.001 and periodic boundary conditions were employed, and electrostatic focusing was turned off due to the presence of the membrane. The water dielectric constant was set to 80.0, and the protein dielectric constant was set to 20. The membrane was modeled as a continuum slab with a thickness of 40 Å and a relative dielectric constant of 7.0. The dielectric boundaries between protein, membrane, and solvent were defined with the classical solvent-excluded surface model using radii for the solvent and membrane probe of 1.4 Å and 2.7 Å, respectively, and using an automatic pore searching algorithm (72).

### Mammalian cell culture

Chinese hamster ovary cells (CHO-K1, CRL 9618, American Type Culture Collection, Manassas VA, USA) were grown in F-12 nutrient medium (GIBCO/Invitrogen, San Diego, CA, USA) supplemented with 10% fetal bovine serum (ATLANTA Biologicals, Norcross, GA, USA), penicillin (50 units/mL), streptomycin (50 μg/mL) at 37°C in 5% CO_2_. The identity of CHO-K1 cells was certified by American Type Culture Collection using cytochrome C oxidase (COI) assay testing. Cells were negative for mycoplasma contamination and are regularly tested using the MycoAlert PLUS Mycoplasma Detection Kit (Lonza, Rockville, MD, USA). Unless stated otherwise, all tissue culture media was obtained from Life Technologies, Inc. (Grand Island, NY, USA). CHO-K1 cells constitutively expressing human KCNE1 (designated CHO-KCNE1 cells) were generated as previously described (74) using the FLP-in™ system (Thermo Fisher Scientific, Waltham, MA, USA) and maintained under selection with hygromycin B (600 μg/mL).

### Electroporation

Plasmids encoding WT or mutant KCNQ1 channels were transiently transfected by electroporation using the Maxcyte STX system (MaxCyte Inc., Gaithersburg, MD, USA). CHO-E1 cells grown to 70-80% confluence were harvested using 0.25% trypsin. A 500 µl aliquot of cell suspension was then used to determine cell number and viability using an automated cell counter (ViCell, Beckman Coulter). Remaining cells were collected by gentle centrifugation (160×g, 4 minutes), washed with 5 ml electroporation buffer (EBR100, MaxCyte Inc.), and re-suspended in electroporation buffer at a density of 10^8^ viable cells/ml. Each electroporation was performed using 100 µl of cell suspension.

CHO-E1 cells were electroporated with 15 µg of WT or mutant KCNQ1 plasmid DNA. The DNA-cell suspension mix was transferred to an OC-100 processing assembly (MaxCyte Inc.) and electroporated using the preset CHO-PE protocol. Immediately after electroporation, 10 µl of DNase I (Sigma-Aldrich, St. Louis, MO, USA) was added to the DNA-cell suspension and the entire mixture was transferred to a 35 mm tissue culture dish for a 30 min incubation at 37°C in 5% CO_2_. Following incubation, cells were gently re-suspended in culture media, transferred to a T75 tissue culture flask and grown for 48 hours at 37°C in 5% CO_2._ Following incubation, cells were harvested, counted, transfection efficiency determined by flow cytometry, and then frozen in 1 ml aliquots at 1.5×10^6^ viable cells/ml in liquid N_2_ until used in experiments.

### Electrophysiology

The day before automated patch clamp recordings, electroporated cells were thawed, plated and incubated for 10 hours at 37°C in 5% CO_2_. The cells were then grown overnight at 28°C in 5% CO_2_ to increase channel expression at the plasma membrane. Prior to experiments, cells were passaged using 5% trypsin in cell culture media. Cell aliquots (500 µl) were used to determine cell number and viability by automated cell counting, and transfection efficiency by flow cytometry. Cells were then diluted to 200,000 cells/ml with an external bath solution (see below) and allowed to recover 60 minutes at 15°C while shaking on a rotating platform at 200 rpm.

Automated patch clamp experiments were performed using the Syncropatch 768 PE platform (Nanion Technologies, Munich, Germany) equipped with single-hole, 384-well recording chips with medium resistance (2-4 MΩ). Pulse generation and data collection were carried out with PatchController384 V.1.3.0 and DataController384 V1.2.1 software (Nanion Technologies, Munich, Germany). Whole-cell currents were recorded at room temperature in the whole-cell configuration, filtered at 3 kHz and acquired at 10 kHz. The access resistance and apparent membrane capacitance were estimated using built-in protocols. The external bath solution contained: 140 mM NaCl, 4 mM KCl, 2 mM CaCl_2_, 1 mM MgCl_2_, 10 mM HEPES, 5 mM glucose, pH 7.4. The internal solution contained: 60 mM KF, 50 mM KCl, 10 mM NaCl, 10 mM HEPES, 10 mM EGTA, 2 mM ATP-K_2_, pH 7.2. Whole-cell currents were elicited from a holding potential of -80 mV using 2000 ms depolarizing pulses (from -80 mV to +60 mV in +10mV steps, every 10 secs) followed by a 2000 ms step to -30 mV to analyze tail currents and channel deactivation rate. Peak currents were recorded 1990 ms after the start of the voltage pulse, while tail currents were measured 10 ms after changing the membrane potential to -30 mV. Non-specific currents were eliminated by recording whole-cell currents before and after addition of the I_Ks_ blocker JNJ-303 (2 μM). Whole-cell currents were not leak-subtracted. Only JNJ303-sensitive currents and recordings meeting the following criteria were used in data analysis: seal resistance ≥ 0.5 GΩ, series resistance ≤ 20 MΩ, capacitance ≥ 1 pF, voltage-clamp stability (defined as the standard error for the baseline current measured at the holding potential for all test pulses being <10% of the mean baseline current). The voltage-dependence of activation was calculated by fitting the normalized G-V curves with a Boltzmann function (tail currents measured at -30 mV). Time constants of channel deactivation were determined by fitting tail currents pre-activated with the +60 mV pulse to a single exponential function. The effect of ML277 on KCNQ1 WT and mutants was determined by measuring currents recorded from cells pre-incubated with 5 µM of ML277 in bath solution for 30 min and compared to currents recorded from cells exposed to vehicle only.

### Electrophysiology data analysis

Data were collected for each experimental condition from at least three transfections and analyzed and plotted using DataController384 V1.8 (Nanion Technologies, Munich, Germany), Clampfit V10.4 (Molecular Devices Corp.), Excel (Microsoft Office 2013, Microsoft), SigmaPlot 2000 (Systat Software, Inc., San Jose, CA, USA) and Prism 8 (GraphPad Software, San Diego, CA)) software. Whole-cell currents were normalized for membrane capacitance and results expressed as mean ± SEM. Additional custom semi-automated data handling routines were used for rapid analysis of current density and voltage-dependence of activation (75). The voltage-dependence of activation was determined only for cells with mean current density greater than the background current amplitude.

## Acknowledgements

This work was supported by NIH grants R01 HL122010 and R01 GM080403. MW was supported by NIH T32 GM008320. The authors acknowledge the computational resources provided by the Advanced Computing Center for Research and Education (ACCRE) at Vanderbilt University and the Center for Scientific Computing at Leipzig University. JM further acknowledges the Deutsche Forschungsgemeinschaft (DFG, German Research Foundation) through SFB1423, project number 421152132.

## Data Availability

Structural models of KCNQ1 and KCNQ1-KCNE1 in complex with ML277 are included as supplementary files **S1** and **S2** to this manuscript. Representative snapshots from the MD simulations as well as example simulation input files have been deposited under: https://doi.org/10.5281/zenodo.5643710. All electrophysiology data generated and analyzed during this study are included in the manuscript or the supplementary material. All data needed to evaluate the conclusions in the paper are present in the paper and/or in the supplementary material.

## Author Contributions

Conceptualization, GK, CGV, CRS, ALG, JM; methodology, GK, CGV, MCW, SA, RRD; formal analysis, GK, CGV, MCW, SA, RRD; investigation, GK, CGV, MCW, SA, RRD; data curation, GK, CGV; validation, GK, CGV; visualization, GK, CGV; writing—original draft preparation, GK, CGV; writing—review and editing, GK, CGV, MCW, SA, RRD, CRS, ALG, JM; supervision, GK, CGV, CRS, ALG, JM; project administration, CRS, ALG, JM; funding acquisition, CRS, ALG, JM.

## Supporting Information

**Figure S1:** ML277 starting positions used for ligand docking.

**Figure S2:** Ten largest clusters of ML277 docking poses after first round of ligand docking.

**Figure S3:** Results of ML277 control docking calculations.

**Figure S4:** Superposition of KCNQ1 with KCNQ2 and KCNQ4 in region of ML277 binding pocket.

**Figure S5:** Whole-cell currents recorded from WT and mutant KCNQ1 channels under control conditions.

**Figure S6:** Whole-cell currents recorded from WT and mutant KCNQ1 channels incubated with ML277.

**Figure S7:** Chemical structures and atom groupings of ligand cofactors used in MD simulations.

**Figure S8:** Molecular contact between KCNE1-F53 and ML277 observed in MD simulations.

**Figure S9:** Change of information flow in the KCNQ1 channel caused by KCNE1.

**Table S1:** Summary of electrophysiological properties of wild-type and mutant KCNQ1 channels studied in this work under control conditions.

**Table S2:** ML277-induced effects on peak and tail currents, voltage-dependence of activation and deactivation kinetics on KCNQ1 mutants.

**File S1:** PDB file of KCNQ1-ML277 model

**File S2:** PDB file of KCNQ1-KCNE1-ML277 model

