## Supplemental Material for "Molecular simulations reveal a mechanism for enhanced allosteric coupling between voltage-sensor and pore domains in KCNQ1 explaining its activation by ML277"

### Supporting Figures

#### Figure S1: ML277 starting positions used for ligand docking

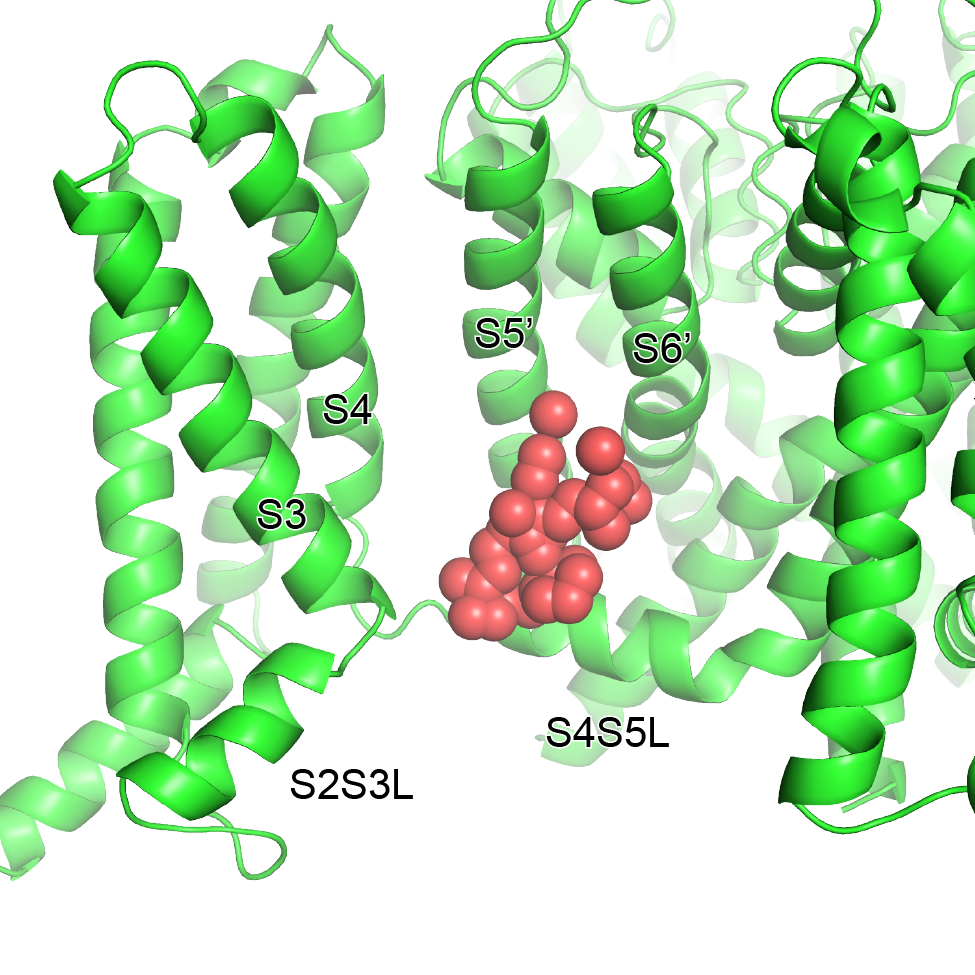

**Figure S1:** ML277 starting positions used for ligand docking are indicated as red spheres.

#### Figure S2: ML277 docking poses after first round of ligand docking

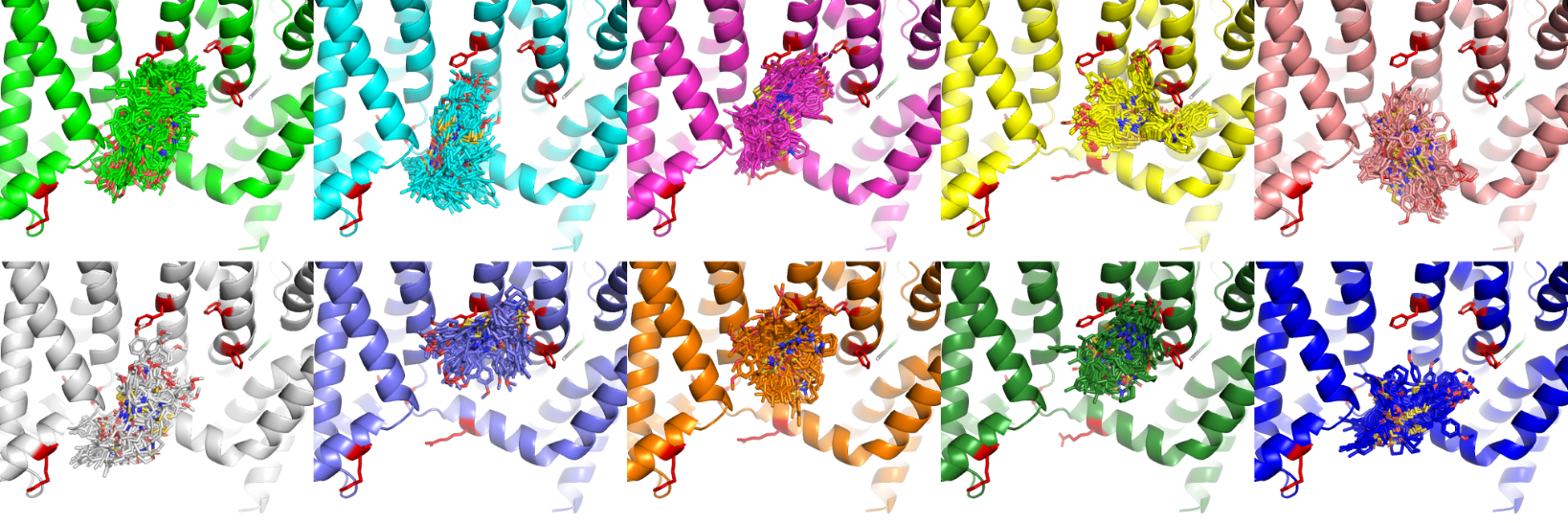

**Figure S2:** Ten largest clusters of ML277 docking poses after first round of ligand docking with RosettaLigand. In some poses, ML277 is above S4S5L and interacts with S5 and S6, while in other poses ML277 is shifted downwards and is between S2S3L, S3, and S4S5L. The best-scoring models of each cluster were used for an additional round of local ligand docking to further refine the ML277 binding poses.

#### Figure S3: ML277 control docking calculations

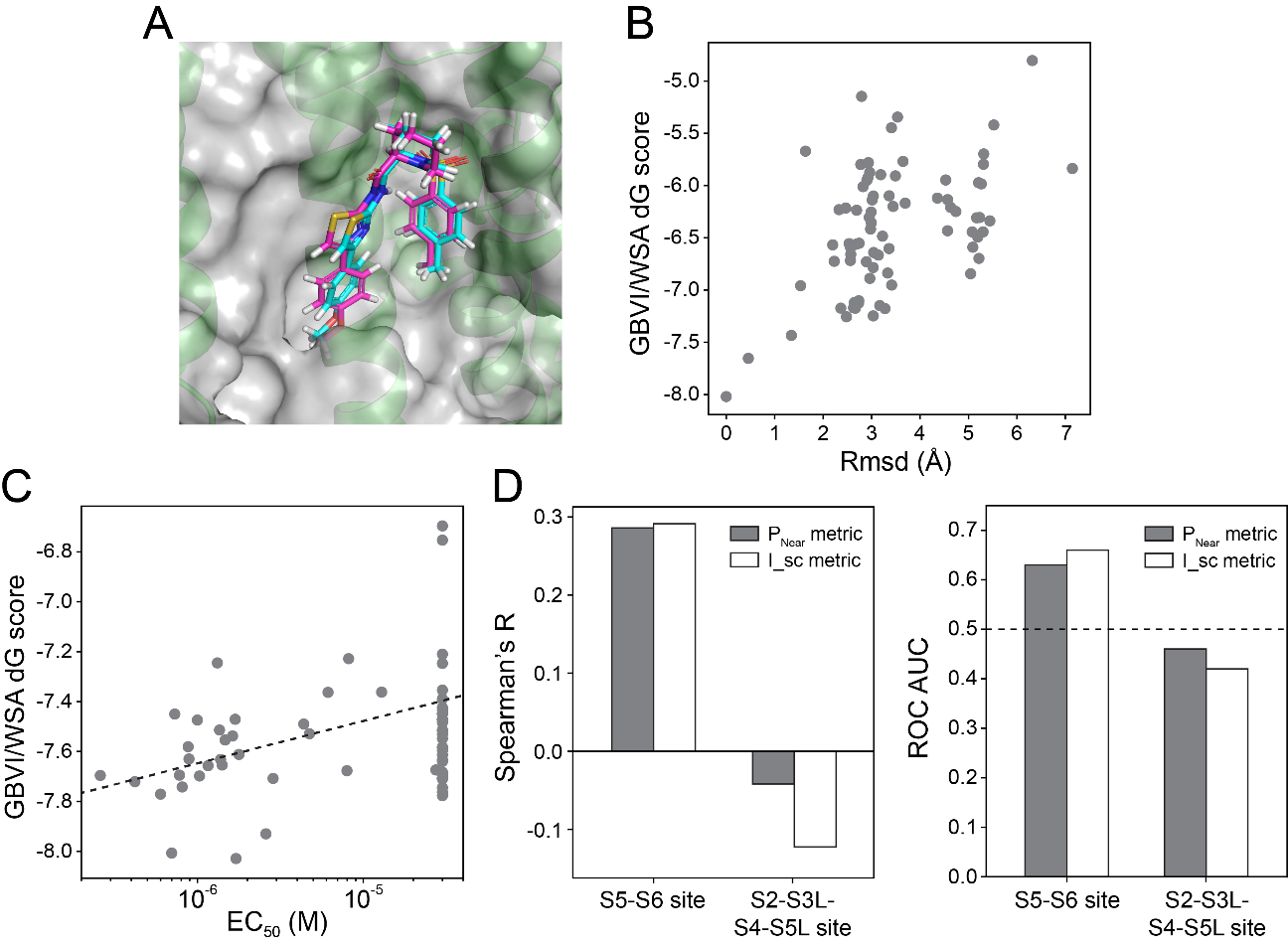

**Figure S3:** Results of ML277 control docking calculations. (**A**) Comparison of best-scoring docking poses of ML277 generated with RosettaLigand (magenta) and MOE software (cyan), respectively. (**B**) Score of ML277 docking models generated with MOE versus ligand-atom root-mean-square distance deviation (Rmsd) relative to best-scoring docking pose. (**C**) GBVI/WSA dG score versus EC_50_ for ML227 and 62 analogs of ML227. The dashed line shows the best linear regression fit to the data (Spearman’s R = 0.3, p = 0.017). (**D**) Spearman’s R and area under the ROC curve obtained for the comparison of Rosetta P_Near_ metric and interface score (I_sc) with EC_50_ data for ML277 and 62 analogs of ML277. The degree of correlation is compared for docking calculations started either from a site close to S5+S6 in the middle of the membrane or a site close to S2-S3L and S4-S5L.

#### Figure S4: Superposition of KCNQ1 with KCNQ2 and KCNQ4 in region of ML277 binding pocket

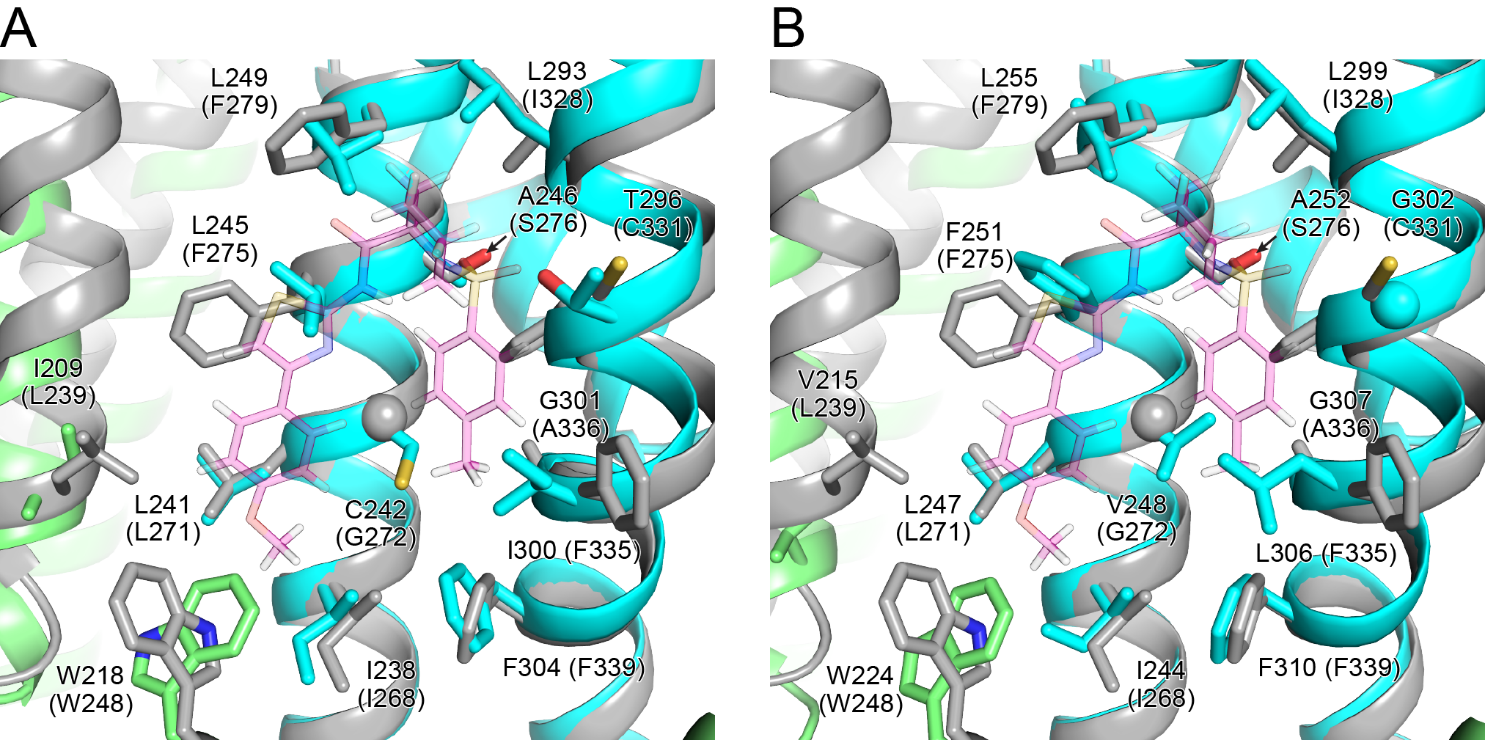

**Figure S4:** Amino acid residues lining the predicted ML277 binding pocket in KCNQ1 have different identity in KCNQ2 and KCNQ4 channels. Superimposition of KCNQ1 with (**A**) KCNQ2 and (**B**) KCNQ4 structures in the ML277 binding pocket. KCNQ2 and KCNQ4 are colored green and cyan. KCNQ1 is colored gray and KCNQ1 residue numbers are shown in parentheses.

#### Figure S5: Whole-cell currents recorded from WT and mutant KCNQ1 channels under control conditions.

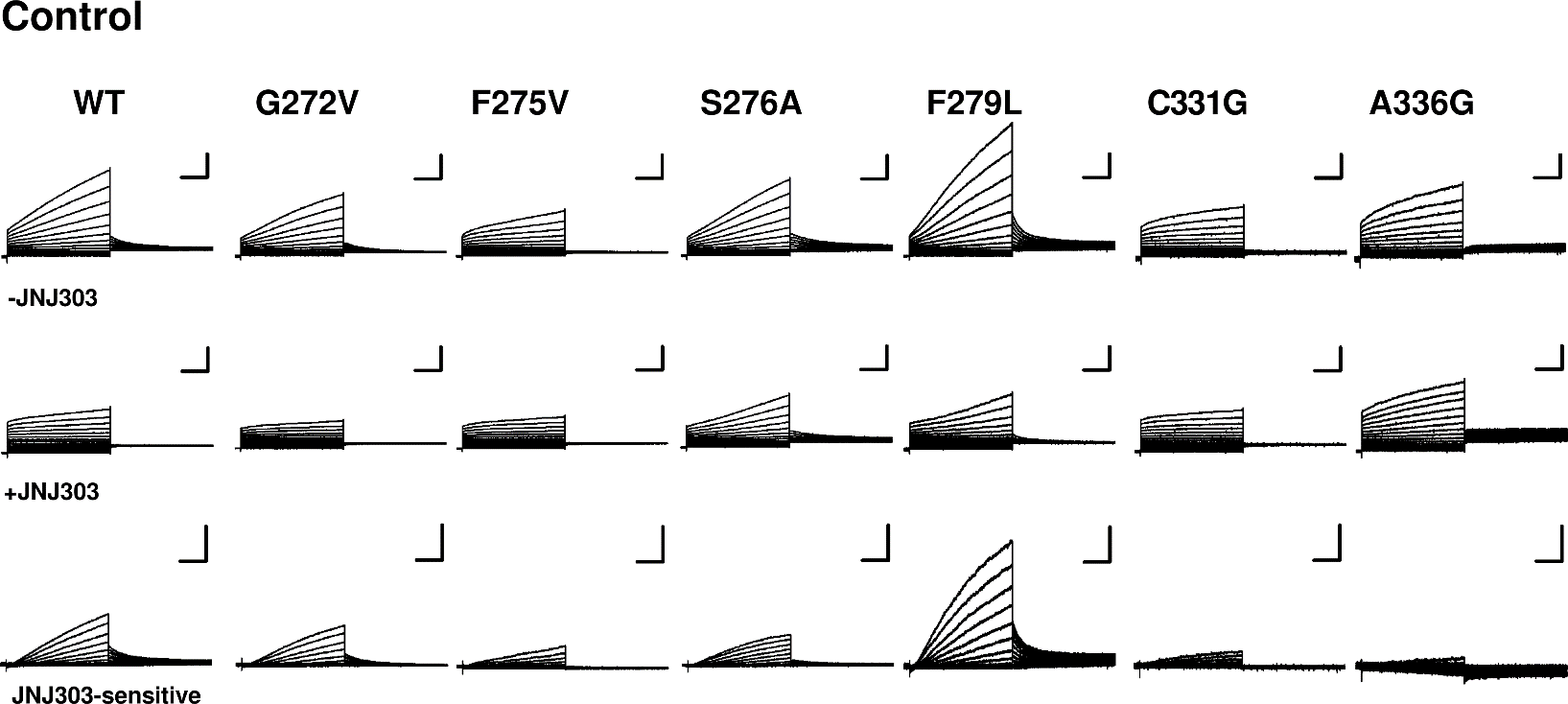

**Figure S5:** Whole-cell currents recorded from WT and mutant KCNQ1 channels under control conditions. Average whole-cell currents normalized by membrane capacitance recorded from CHO-E1 cells transiently expressing WT or mutant KCNQ1 channels exposed to vehicle (control) before and after JNJ303 addition. Vertical scale = 25 pA/pF, horizontal scale bar = 500 ms.

#### Figure S6: Whole-cell currents recorded from WT and mutant KCNQ1 channels incubated with ML277.

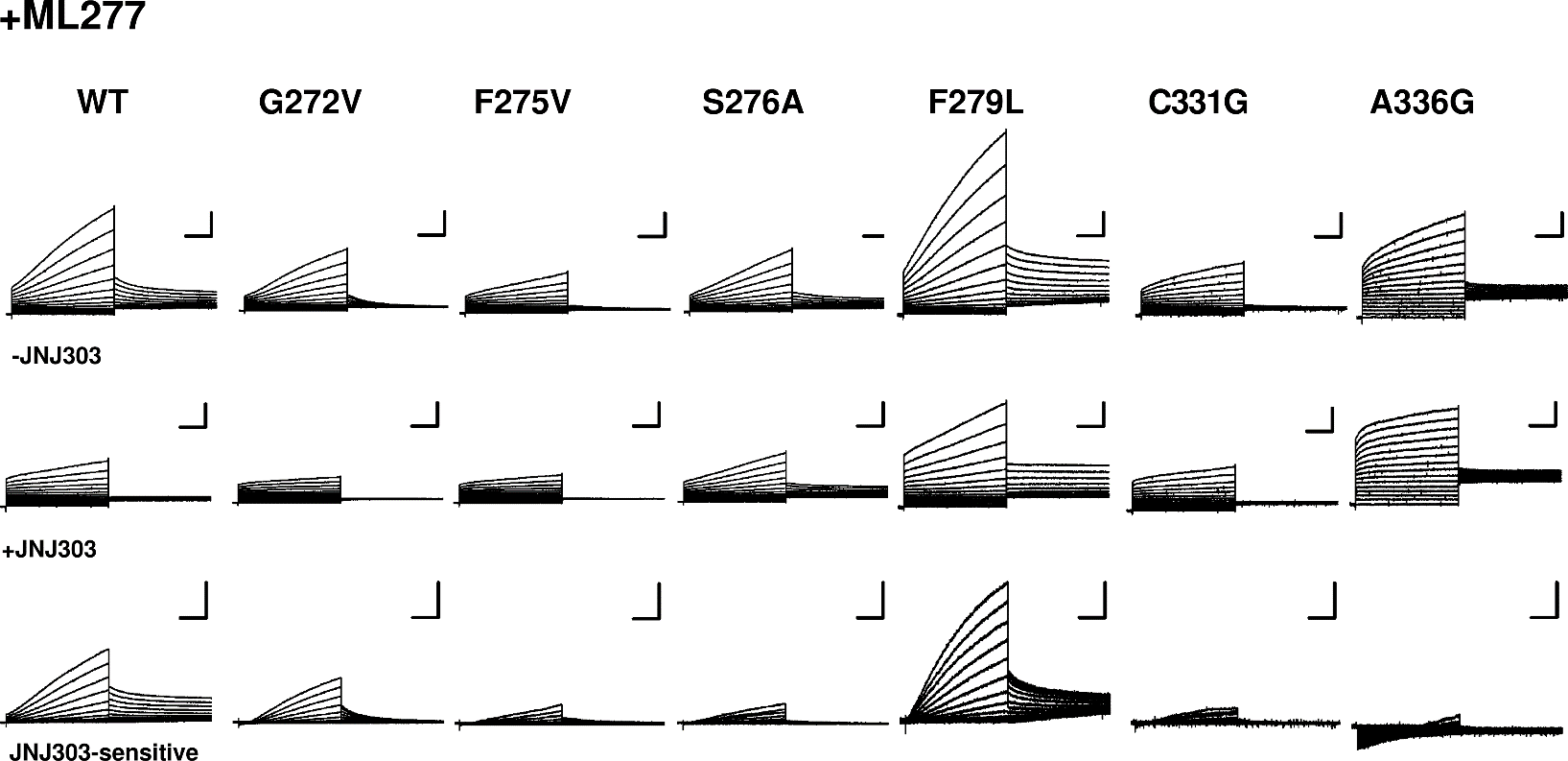

**Figure S6:** Whole-cell currents recorded from WT and mutant KCNQ1 channels incubated with ML277. Average whole-cell currents normalized by membrane capacitance recorded from CHO-E1 cells transiently expressing WT or mutant KCNQ1 channels exposed to ML277 before and after JNN303 addition. Vertical scale = 25 pA/pF, horizontal scale bar = 500 ms,

#### Figure S7: Cofactor atom grouping used for network building

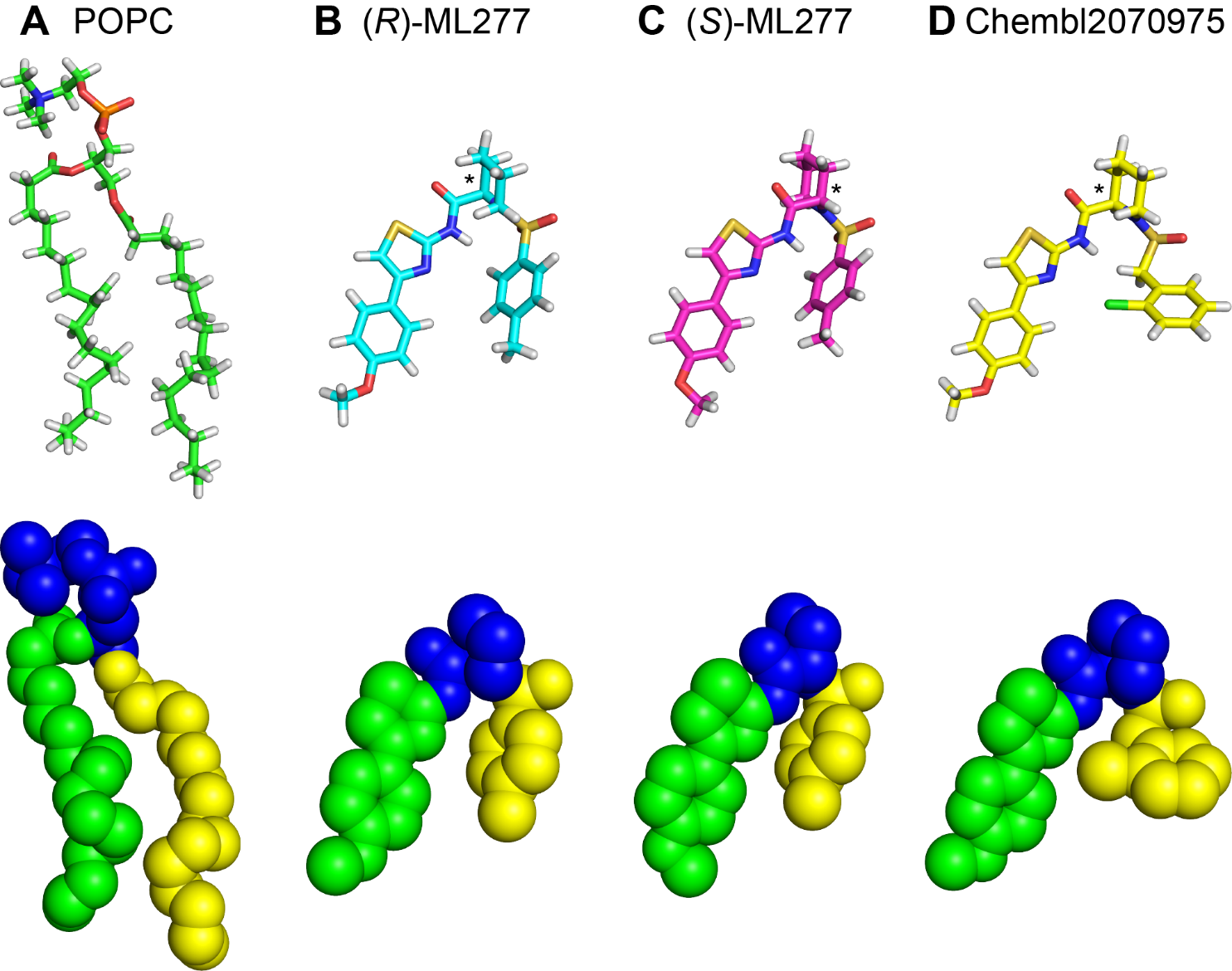

**Figure S7:** Chemical structures and atom groupings of ligand cofactors used in MD simulations in this study. (**A**) POPC, (**B**) (*R*)-ML277, (**C**) (*S*)-ML277, (**D**) CHEMBL2070975. The grouping of atoms into domains is shown with different colors. Domains were assigned nodes in the MD network.

#### Figure S8: Contact between KCNE1 and ML277 observed in MD simulations

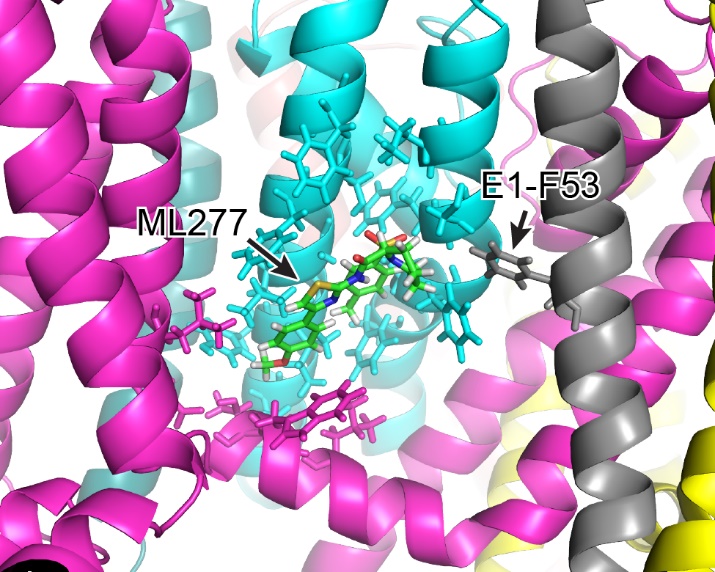

**Figure S8:** A molecular contact between KCNE1-F53 and ML277 was observed in MD simulations.

#### Figure S9: Change of information flow in the KCNQ1 channel caused by KCNE1

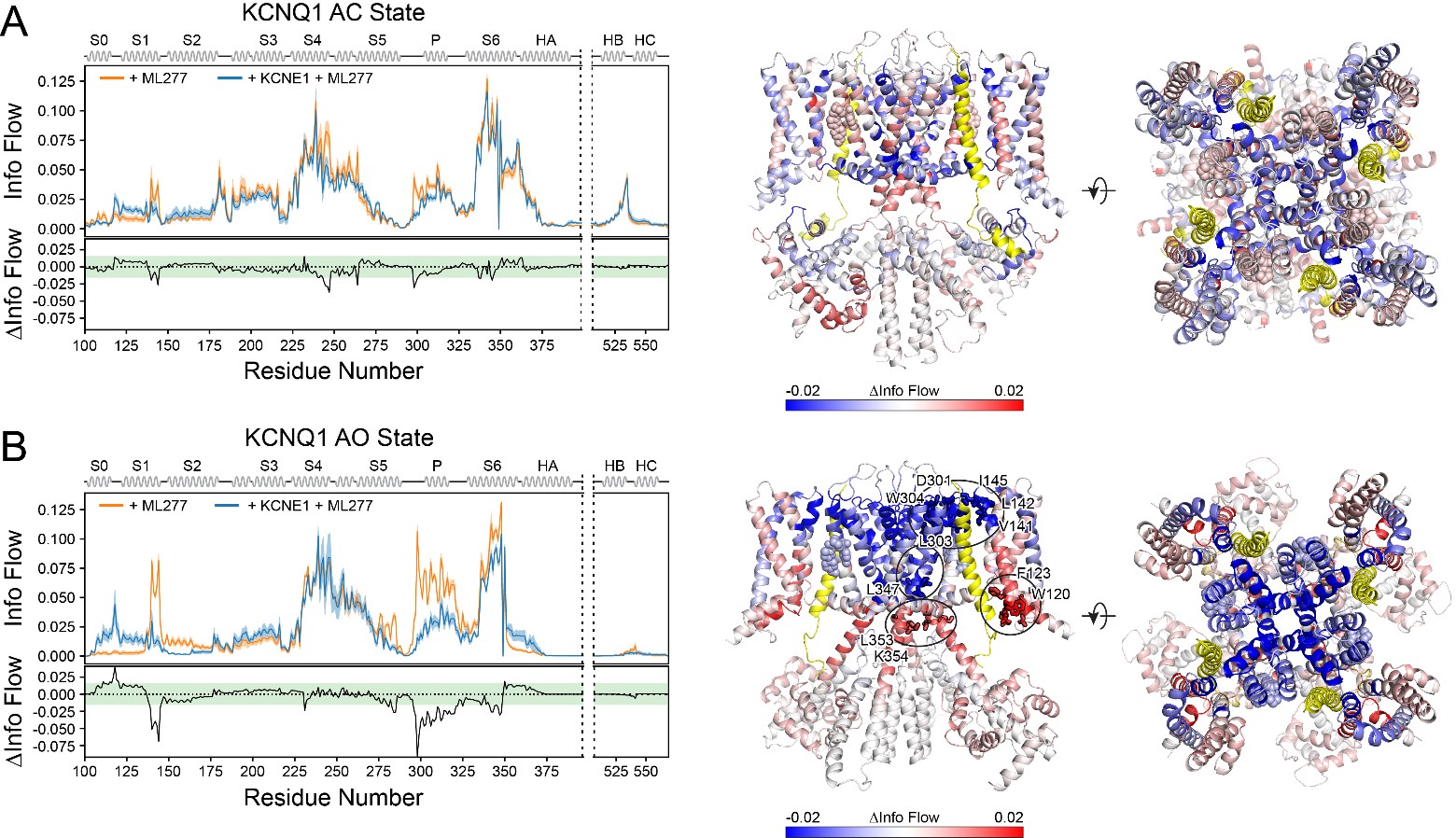

**Figure S9:** KCNE1 changes allosteric signaling mostly in the KCNQ1 AO state structure. (**A**) Left: Information flow through the KCNQ1 AC structure with either ML277 or ML277+KCNE1 bound and the difference in information flow between simulations with and without KCNE1 (Δ Info Flow) are plotted along the KCNQ1 residue sequence. Right: The change of information flow induced by KCNE1 (capped at ±0.02) is projected onto the KCNQ1 AC structure. The VSDs, CaMs, ML277, and KCNE1 molecules in the front and back are omitted in the left image with the viewing direction parallel to the membrane plane. KCNE1 is shown in yellow. (**B**) Left: Information flow through the KCNQ1 AO structure in the presence of either ML277 or ML277+KCNE1 and the information flow difference between the two conditions are plotted along the KCNQ1 residue sequence. Right: The change of information flow induced by KCNE1 is mapped on the KCNQ1 AO structure. Regions which become less important for allosteric signaling between VSD and PD in the presence of KCNE1 are indicated by dark blue color. These regions include the P helix and selectivity filter, the extracellular halves of S5 and S6, and the extracellular end of S1. Regions in which information flow is increased by the presence of KCNE1 are colored red and include the lower half of S1, the intracellular end of S6 and the part connecting S6 to helix HA. Hotspot residues showing the largest changes in information flow are displayed as sticks and labeled. KCNE1 is colored yellow.

### Supporting Tables

|  | **WT** | | | **G272V** | | | **F275V** | | | **S276V** | | | **F279L** | | | **C331G** | | | **A336G** | | |
| --- | --- | --- | --- | --- | --- | --- | --- | --- | --- | --- | --- | --- | --- | --- | --- | --- | --- | --- | --- | --- | --- |
|  | **avg** | **sem** | **n** | **avg** | **sem** | **n** | **avg** | **sem** | **n** | **avg** | **sem** | **n** | **avg** | **sem** | **n** | **avg** | **sem** | **n** | **avg** | **sem** | **n** |
| **Ipeak, pA/pF** | 52.14 | 3.47 | 156 | 37.66* | 3.94 | 57 | 16.39* | 2.19 | 47 | 22.85* | 2.80 | 69 | 81.88* | 8.11 | 23 | 10.74* | 1.67 | 42 | 13.1* | 7.11 | 12 |
| **Itail, pA/pF** | 18.10 | 1.29 | 156 | 10.53* | 1.30 | 57 | 0.14* | 0.55 | 44 | 5.90* | 1.30 | 69 | 30.18* | 3.24 | 23 | -0.53* | 0.64 | 42 | -2.48* | 4.14 | 13 |
| **V½, mV** | 29.54 | 0.58 | 96 | 30.16 | 1.13 | 37 | 38.66* | 0.95 | 10 | 23.45* | 1.75 | 35 | 24.45* | 1.55 | 24 |  | nd |  |  | nd |  |
| **Deac τ, ms** | 343.0 | 1.6 | 156 | 405.9* | 1.9 | 57 | 598.5* | 25.9 | 47 | 502.1* | 6.17 | 69 | 220.1* | 1.3 | 23 | 261.8* | 17.8 | 42 | 310.3 | 9.3 | 12 |

#### Table S1**:** Summary of electrophysiological properties of wild-type and mutant KCNQ1 channels studied in this work under control conditions.

* p ≤ 0.01

|  | **WT** | | | **G272V** | | | **F275V** | | | **S276V** | | | **F279L** | | | **C331G** | | | **A336G** | | |
| --- | --- | --- | --- | --- | --- | --- | --- | --- | --- | --- | --- | --- | --- | --- | --- | --- | --- | --- | --- | --- | --- |
|  | **avg** | **sem** | **n** | **avg** | **sem** | **n** | **avg** | **sem** | **n** | **avg** | **sem** | **n** | **avg** | **sem** | **n** | **avg** | **sem** | **n** | **avg** | **sem** | **n** |
| **Ipeak, %** | 128.5* | 8.6 | 156 | 109.7 | 11.0 | 63 | 101.7 | 12.8 | 48 | 69.5 | 15.6 | 62 | 106.9 | 15.6 | 26 | 84.6 | 16.9 | 42 | 12.5 | 39.2 | 12 |
| **Itail, %** | 174.5* | 11.7 | 156 | 141.2 | 14.9 | 63 | 822.4* | 201.2 | 48 | 10.5* | 21.9 | 48 | 107.4 | 16.7 | 26 | 193.0 | 62.8 | 42 | 96.6 | 32.3 | 12 |
| **△V½, mV** | -1.80 | 0.77 | 121 | -0.05 | 0.85 | 49 | 1.51 | 1.60 | 13 | 0.71 | 2.93 | 20 | -8.19* | 2.72 | 20 |  | nd |  |  | nd |  |
| **Deac τ, M/C** | 1.22* | 0.00 | 156 | 0.95* | 0.01 | 63 | 0.84 | 0.05 | 48 | 1.76* | 0.01 | 62 | 2.09* | 0.00 | 26 | 1.47* | 0.05 | 42 | 1.00 | 0.03 | 12 |

#### Table S2**:** ML277-induced effects on peak and tail currents, voltage-dependence of activation and deactivation kinetics on KCNQ1 mutants.

* p ≤ 0.01
